# Mechanical control of histone serotonylation initiates neural crest migration *in vivo*

**DOI:** 10.1101/2025.10.21.683626

**Authors:** Joana E. Saraiva, Artemis G. Korovesi, Xiaoran Wei, Jaime A. Espina, Miguel Ribeiro, Ian Maze, Elias H. Barriga

**Affiliations:** Biophysical Mechanisms of Morphogenesis Lab, Cluster of Excellence Physics of Life (PoL), TU Dresden, Dresden, Germany; Mechanisms of Morphogenesis Lab, Gulbenkian Institute of Science (IGC); Oeiras, Portugal; Patterning and Morphogenesis Lab, Gulbenkian Institute of Science (IGC); Oeiras, Portugal; Nash Family Department of Neuroscience, Friedman Brain Institute, Icahn School of Medicine at Mount Sinai, New York, NY, 10029, USA; Department of Pharmacological Sciences, Icahn School of Medicine at Mount Sinai, New York, NY, 10029, USA; Howard Hughes Medical Institute, Icahn School of Medicine at Mount Sinai, New York, NY, 10029, USA

**Keywords:** Chromatin, tissue mechanics, cell states transitions, Tgm2, Histone serotonylation, collective cell migration, neural crest, *Xenopus*

## Abstract

Collective cell migration (CCM) is pivotal in several biological contexts, and posttranslational modifications of histones are essential to initiate this process^1–3^. Here, we show that a recently discovered chromatin mark, termed histone serotonylation^4^, is involved in the collective migration of cranial neural crest cells – an embryonic multipotent stem cell population^5^. Our *in vivo* data reveal that histone serotonylation appears in neural crest cells just before they start migrating and that its occurrence is essential to initiate their CCM. Surprisingly, we found that stiffening of the neural crest migratory substrate, the mesoderm, induces histone serotonylation by promoting nuclear translocation of transglutaminase 2 (Tgm2), the enzyme responsible for adding serotonin to histones^4^. Moreover, mechanical and molecular perturbations demonstrate that mechanical shuttling of Tgm2 into the nucleus, with concomitant increases in histone serotonylation, are both required and sufficient to allow CCM *in vivo*. Furthermore, integrated chromatin immunoprecipitation and RNA sequencing analyses uncover a transcriptional module, which is enabled by histone serotonylation in response to mesoderm stiffening. Altogether, our results provide *in vivo* evidence showing that tissue stiffening leads to increased levels of histone serotonylation to reinforce permissive patterns of gene expression, supporting the switch from non-migratory to migratory cell states.

## Main Text

A number of chromatin posttranslational modifications (PTMs), such as histone methylation and acetylation, are involved in the initiation of collective cell migration (CCM) of neural crest cells and other cell-types^1–3^. These PTMs function primarily to fine-tune the chromatin landscape and binding to transcriptional regulators in order to allow for appropriate patterns of gene expression that drive cell fate decisions and plasticity^4,6–8^. One such mark, histone H3 lysine (K) 4 trimethylation (H3K4me3), has been demonstrated to be required for neural crest CCM^9^ and plays critical roles in the reinforcement of permissive gene expression. Emerging evidence suggests that the maintenance of H3K4me3 is influenced by adjacent H3 glutamine (Q) 5 serotonylation^4^– a recently discovered chromatin PTM which is *written*, *erased* and *exchanged* by the transglutaminase 2 (TGM2) enzyme^10,11^. Mechanistically, serotonylation of histone H3 in Q5 enhances the binding of H3K4me3 *reader* proteins (e.g., TFIID complex) and inhibits interactions with KDM5 and LSD1 demethylases that remove H3K4 methylation^4,12^. Given the importance of both H3K4me3 and serotonin (5-HT) in the development of neural crest cells^13–16^, we investigated whether histone serotonylation is involved in the collective migration of neural crest cells.

We first analyzed the occurrence of histone serotonylation in non-migratory, pre-migratory and migratory neural crest cells *in vivo* (**Fig. 1a-c**). Temporal profiling revealed a robust increase in histone serotonylation levels when neural crest cells transit from non-migratory to pre-migratory stages, which is sustained during migration (**Fig. 1b, c**). Next, we investigated whether this increase in histone serotonylation plays a role in neural crest CCM. For this, we used a previously described catalytically dead mutant form of histone H3^4^ (**Supplementary Fig. 1a, b**), which owing to the replacement of H3Q5 with an alanine (A; H3Q5A) prevents histone serotonylation. Microinjection of a wildtype form of H3 (WT H3) did not affect histone serotonylation, but H3Q5A microinjection reduced neural crest histone serotonylation both *in vivo* (**Fig. 1d-f**) and *ex vivo* (**Supplementary Fig. 1c, d**). This confirms the effectiveness of this construct to affect histone serotonylation as well as the specificity of the antibody to detect this mark. Next, *in situ* hybridization analyses against specific neural crest migration marker (*sox8*) revealed that H3Q5A-treated cells failed to initiate their collective movement when compared to animals treated with a WT H3, for which stereotypical migratory streams were evident (**Fig. 1g-i**). Our *ex vivo* migration assays also demonstrated that isolated H3Q5A-treated neural crest clusters failed to spread and migrate (**Supplementary Fig. 1e-g**), confirming that the impact on CCM is not due to the potential requirement of histone serotonylation in other neural tissues. Together, these observations indicated that histone serotonylation already occurs in the pre-migratory neural crest and that it is required for the onset of its CCM *in vivo*.

**Figure 1.**
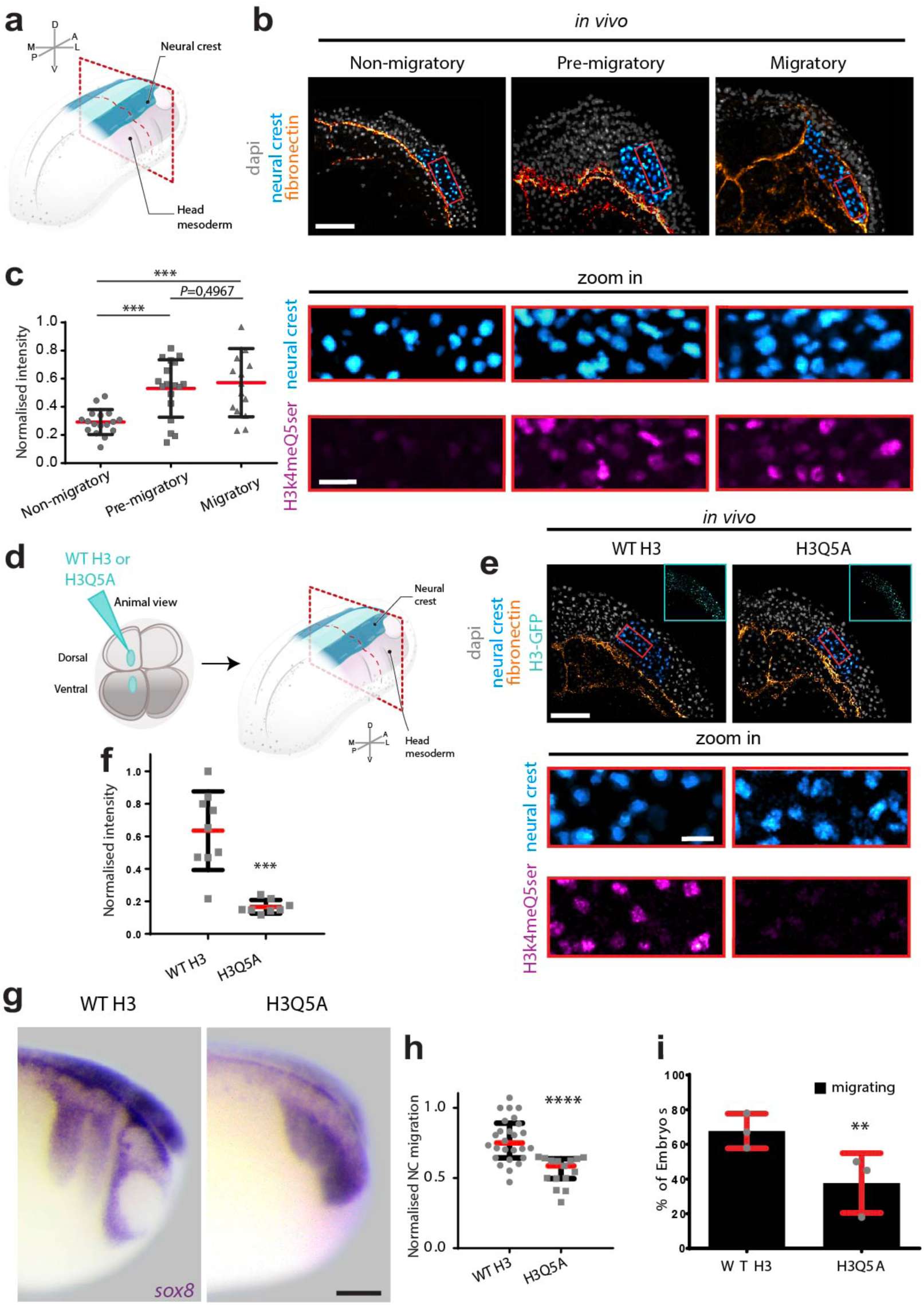
Histone serotonylation is required for neural crest collective cell migration. **a**, Diagram of a *X. laevis* embryo in a pre-migratory stage, the position of the cephalic neural crest and the mesoderm are indicated, red dashed line represents the transverse plane of cryosections (ML, mediolateral; AP, anteroposterior; DV, dorsoventral). **b,** Representative confocal projections of transverse cryosections showing neural crest at non-migratory (stage 13), pre-migratory (stage 18) and migratory stages (stage 21); scale bar, 200 µm; red square indicates the region of interest shown in the lower panels (zoom-in with scale bar, 40 µm); conditions as indicated. **c,** Normalised histone serotonylation fluorescence intensity (details in Methods); red lines represent mean and error bars the standard deviation; two-tailed t test with Welch’s correction \*\*\**P*<0.0001, *P*<0.0005, n_(non-migratory)_ = 17 embryos; n_(pre-migratory)_ = 18 embryos; n_(migratory)_ = 15 embryos. **d,** Schematic shows targeted injections (details in Methods) and red dashed line depicts the plane of transverse cryosection. **e,** Representative confocal projections (top-right inset shows construct expression); scale bar, 200 µm; red square indicates the region of interest shown in the lower panel (zoom-in; scale bar, 40 µm); conditions as indicated. **f,** Normalised histone serotonylation fluorescence intensity; red lines represent mean and error bars the standard deviation; two-tailed t test with Welch’s correction *** *P<*0.0004, n_WT H3Q5_=9 embryos; n_(H3Q5A)_ =9 embryos. **g,** *In situ* hybridizations, lateral views of *sox8* hybridized embryos; scale bar, 150 μm; conditions as indicated. **h,** Normalized neural crest migrated distances; spread of data from the indicated conditions is shown, red lines represent median and whiskers represent interquartile ranges, two-tailed Mann Whitney test, **** *P* < 0.0001, n_(WT H3)_ = 26 embryos; n_(H3Q5A)_ = 15 embryos (Stage 23). **i,** Percentage of embryos displaying neural crest migration; Histograms show mean, error bars represent standard deviation; two-tailed fisher test ** *P*=0,00423. Panels in b, e and g are representative examples of at least three independent experiments, CI = 95%.

To gain further insights into the mechanism controlling histone serotonylation in neural crest cells, we next investigated Tgm2, the *bona fide* H3 serotonylase. We first confirmed that *tgm2* is expressed in neural crest cells by *in silico* analysis of neural crest-specific RNA libraries (**Supplementary Fig. 2a, b**). This expression was spatially corroborated by *in situ* hybridization, which revealed the expression of *tgm2* in the neural crest, the eye and neural tube (**Methods**; **Supplementary Fig. 2c, d**). Using immunofluorescence, we additionally detected Tgm2 protein expression in the nuclei of migratory neural crest cells, where it is expected to promote histone serotonylation (**Supplementary Fig. 2e**). Note that this signal was reduced upon Tgm2 knockdown, and increased upon overexpression of full-length *Xenopus laevis* Tgm2 (**Supplementary Fig. 2e**), confirming that this antibody detects *Xenopus* Tgm2. Thus, we analyzed the temporal dynamics of Tgm2 nuclear localization from non-migratory to migratory stages and surprisingly we found that Tgm2 appears in the nuclei of neural crest cells just before the onset of neural crest CCM **(Fig. 2a-c)**. Therefore, we next assessed the requirement of nuclear Tgm2 for neural crest histone serotonylation and the onset of CCM. To dissect nuclear roles for Tgm2 from its cytosolic functions, which have been described in other cell-types^17,18^, a NLS-tagged mutant (C277A) form of Tgm2 previously shown to prevent histone serotonylation was used^10^ referred hereafter as NLS-mutant Tgm2 (**Fig. 2d; Methods**). We first confirmed that NLS-mutant Tgm2 exclusively localizes to neural crest nuclei (**Fig. 2e**) and that its injection prevents histone serotonylation in comparison to injection of a NLS-tagged wildtype form of Tgm2 (**Fig. 2e, f**). We observed that injection of NLS-mutant Tgm2 inhibited neural crest CCM both *in vivo* (**Fig. 2g-i**) and *ex vivo* (**Supplementary Fig. 2f-h)–** indicating the requirement of nuclear Tgm2 for the onset of neural crest CCM, and providing complementary evidence for the requirement of histone serotonylation in this process.

**Figure 2.**
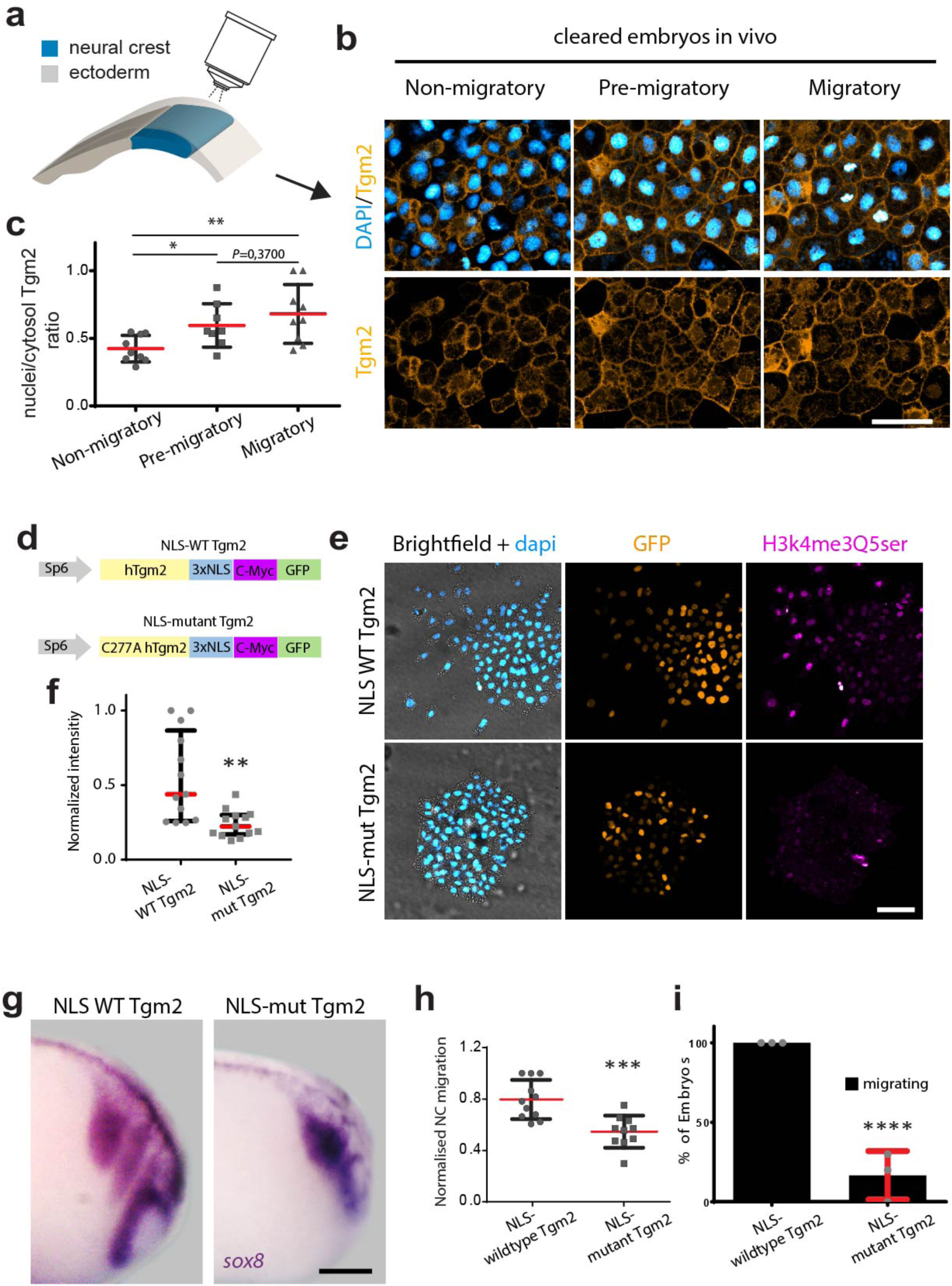
Nuclear Tgm2 is required for histone serotonylation and CCM. **a**, Schematic of the method used to image cleared and flat-mounted embryos (details in Methods). **b,** Tgm2-GFP subcellular localisation at non-(stage 13), pre- (stage 17) and migratory stages (stage 21) is shown in orange; nuclei are shown in cyan; scale bar, 50 μm; conditions as indicated. **c,** Nuclear/cytosolic ratio of Tgm2-GFP fluorescence intensity; red lines represent mean and error bars standard deviation; two-tailed t test with Welch’s correction, \*\**P*=0,0080, n_(non-migratory)_ = 9 embryos; *n*_(pre-migratory)_ =9 embryos; *n*_(migratory)_ = 9 embryos. **d,** Schematic showing Tgm2 constructs used to express NLS-tagged wild-type and NLS-tagged mutant (C277A) forms of Tgm2. **e,** Immunostaining against histone serotonylation in neural crest cells (magenta) after injection of NLS-tagged wild type or NLS-tagged mutant (C277A) forms of Tgm2 (orange); nuclei are shown in cyan; scale bar, 50 μm. **f,** Normalised histone serotonylation fluorescence intensity; spread of data from the indicated conditions is shown, red lines represent median and whiskers represent interquartile ranges (two-tailed Mann Whitney test ** *P*=0.0023, n_(NLS-wildtype Tgm2l)_ = 14 explants nuclei; n_(NLS-mutant Tgm2)_ = 14 explants nuclei. **g,** *In situ* hybridisation, lateral views of *sox8* hybridized embryos, scale bar, 150 μm; conditions as indicated. **h,** Normalized neural crest migrated distances, two-tailed t test with Welch’s correction, *** *P* < 0.0017; n_(NLS-wildtype Tgm2)_ = 11 embryos; n_(NLS-mutant Tgm2)_ = 11 embryos (Stage 23). **i,** Percentage of embryos displaying neural crest migration; histograms show mean; error bars represent standard deviation; two-tailed fisher test **** *P*<0,0001. Panels in b, e and g are representative examples of three independent experiments, CI = 95%.

Given its recent discovery, the environmental or epigenetic factors that modulate the occurrence of histone serotonylation remain largely unknown. In *Xenopus,* the onset of neural crest CCM is mechanically triggered by stiffening of the tissue that they use as a migratory substrate, the mesoderm^5^. Considering that mechanical stimuli also instruct chromatin modifications^19^, we next tested the involvement of mesoderm stiffening on histone serotonylation. For this, we generated embryos with softened mesoderm by using a previously validated tool relying on a targeted injection of an active form of myosin phosphatase-1 (ca-Mypt1)^5,20^ **(Supplementary Fig. 3a, b)**. Remarkably, mesoderm softening resulted in a non-autonomous reduction of histone serotonylation within the neural crest (**Fig. 3a-c**). To more directly test roles for substrate mechanics in the regulation of histone serotonylation, we used an *ex vivo* assay relying on the use of fibronectin-coated hydrogels carrying stiffness values similar to those found in the mesoderm at non-migratory (soft) and early migratory (stiff) stages (**Supplementary Fig. 3c-e**)^5^. In this scenario, we found that cells within neural crest clusters plated on stiff hydrogels displayed higher levels of histone serotonylation, similar to those observed at migratory stages *in vivo*. However, exposure to more compliant surfaces was sufficient to reduce the occurrence of this mark to levels equivalent to those reported in non-migratory neural crests (**Fig. 3d, e**). Complementing these findings, we assessed the relevance of mechanosensing for histone serotonylation *in vivo*. Since it is described that vinculin is part of the mechanosensitive machinery used by neural crest cells to sense mesoderm stiffening and collectively migrate^5^, we used a validated mutant form of Vinculin (Vinculin-Cter), which is known to affect the onset of neural crest CCM^5^. Immunofluorescence analysis showed that histone serotonylation levels are reduced upon vinculin depletion, both *ex vivo* (**Fig. 3d, e**) and *in vivo* (**Fig. 3a-c**). Furthermore, Vinculin depletion was sufficient to inhibit nuclear localization of Tgm2-GFP (**Supplementary Fig. 5a, b**), suggesting a requirement of mechanosensitive machineries, and eventually substrate mechanics, for Tgm2 nuclear translocation. To more directly test this, we exposed neural crest cells to soft or stiff hydrogels and analyzed whether mechanical signaling influences endogenous levels of Tgm2. Immunofluorescence analyses revealed that endogenous Tgm2 localizes to the nuclei of neural crest cells plated on stiff hydrogels, but that exposure to non-migratory stiffness values (soft) was sufficient to exclude Tgm2 from the nuclei of these cells (**Fig. 3f, g**). We also found that *tgm2* mRNA levels do not display major changes from non-migratory to migratory stages and are not affected by vinculin depletion **(Supplementary Fig. 5c, d).** Intriguingly, we observed that mechanical signaling does not affect the co-occurrence of H3K4me3 **(Supplementary Fig. 4a-e)** implying a specific regulation of histone serotonylation. Together, these complementary data sets indicate that mesoderm stiffening and mechanical signaling induce Tgm2 nuclear translocation and with that histone serotonylation at the onset of neural crest CCM.

**Figure 3.**
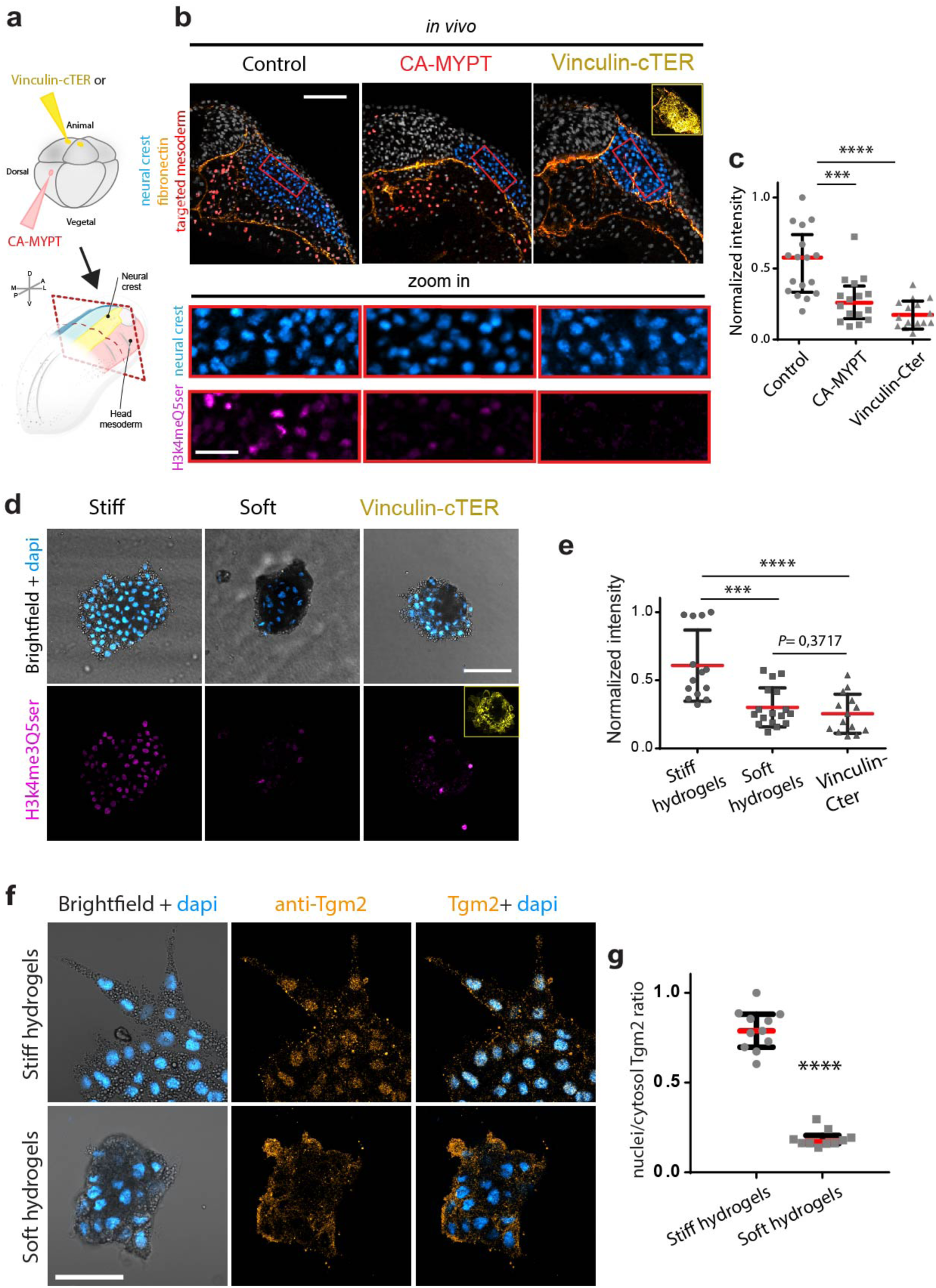
Histone serotonylation and Tgm2 are controlled by substrate mechanics. **a**, Schematic shows targeted injections and red dashed line depicts the plane of transverse cryosection. **b,** Representative confocal projections (top-right inset shows construct expression); scale bar, 200 µm; red square indicates the region of interest shown in the lower panel (zoom-in; scale bar, 40 µm); conditions as indicated. **c,** Normalised histone serotonylation fluorescence intensity; spread of data from the indicated conditions is shown, on CA-MYPT-treated embryos red lines represent median and whiskers represent interquartile ranges (two-tailed Mann Whitney test *** *P<*0.0007, n_(control)_ = 9 embryos; n_(CA-MYPT)_ = 9 embryos; on Vinculin-cTER-treated embryos red lines represent mean and error bars the standard deviation; two-tailed t test with Welch’s correction **** *P*<0.0001; n_(control)_ = 9 embryos; n_(vinculin-cTER)_ = 9 embryos. **d,** Immunostaining against histone serotonylation (magenta) in wild type neural crest clusters platted on stiff or soft hydrogels; and on vinculin-treated clusters (vinculin-cTER) exposed to stiff gels (yellow top-right inset shows correct injection); nuclei are shown in cyan; scale bar, 100 μm; conditions as indicated. **e,** Normalised histone serotonylation fluorescence intensity; red lines represent mean and error bars the standard deviation; two-tailed t-test, \*\*\**P* < 0.0002 and **** *P* < 0.0001; n_(stiff)_= 14 explants; *n*_(soft)_= 17 explants; *n*_(vinculin-Cter)_ = 14 explants. **f,** Immunostaining for endogenous Tgm2 (orange) in wild type neural crest clusters platted on stiff and soft substrates; nuclei are shown in cyan (scale bar, 50 μm); conditions as indicated. **g,** Nuclear/cytosolic ratio of endogenous Tgm2 intensity; spread of data from the indicated conditions is shown, red lines represent median and whiskers represent interquartile ranges, two-tailed Mann–Whitney U-test, \*\*\*\**P* < 0.0001, n_(stiff)_= 11 explants; *n*_(soft)_= 10 explants Panels in b, d and f are representative examples of three independent experiments, CI = 95%.

To confirm the relevance and sufficiency of Tgm2 nuclear shuttling for histone serotonylation and CCM, we overexpressed NLS-tagged wildtype Tgm2 at non-migratory stages where histone serotonylation and CCM are not yet occurring (**Fig. 4a**). Strikingly, this treatment was sufficient to prematurely trigger both histone serotonylation (**Supplementary Fig. 6a, b**) and the onset of neural crest CCM *in vivo* (**Fig. 4b, c**). Premature neural crest migration is also reported to appear upon early stiffening of the mesoderm^5^. Thus, we used this scenario to directly test whether Tgm2 nuclear shuttling and histone serotonylation are force dependent *in vivo*. For this, we resorted to a method relying on the use of an atomic force microscope (AFM) to directly apply extrinsic stress to the migratory path of neural crest cells^5,21^ (**Supplementary Fig. 6c, d; Methods**). As previously reported^5^, early mesoderm stiffening triggered premature neural crest CCM in wildtype embryos (**Supplementary Fig. 6e**), as well as premature Tgm2 nuclear translocation and an early increase of histone serotonylation signal **(Fig. 4d-h**). Furthermore, premature neural crest migration was inhibited in animals treated with NLS-mutant Tgm2 or with mutant H3 (H3Q5A) (**Fig. 4i, j**). Together, these results indicated that mesoderm stiffening is both necessary and sufficient to promote Tgm2 nuclear shuttling, allowing for histone serotonylation and the onset of neural crest CCM *in vivo*.

**Figure 4.**
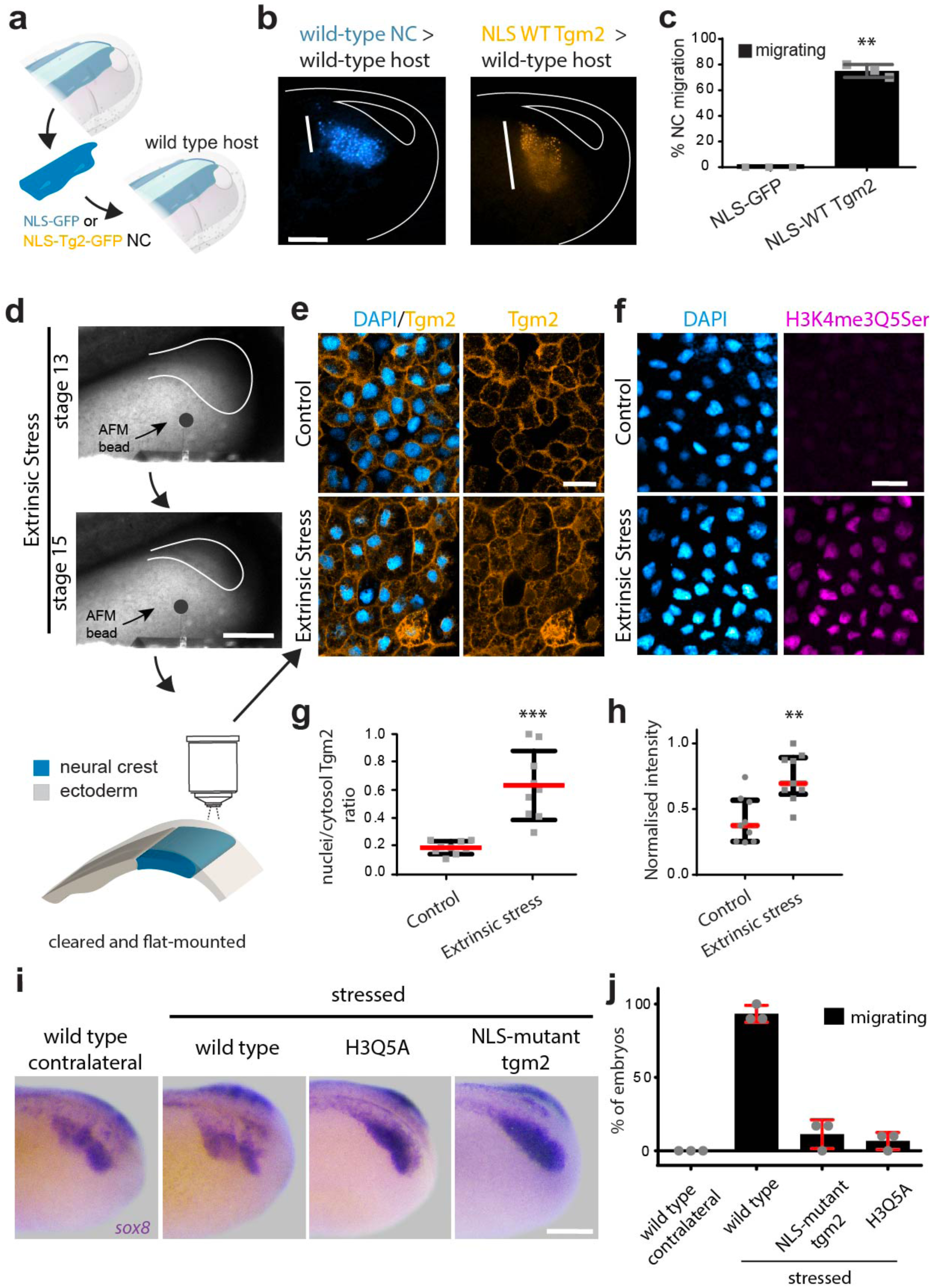
Mesoderm stiffening triggers CCM via Tgm2 nuclear shuttling and histone serotonylation. **a**, Diagram showing graft experiments in which neural crest from NLS-GFP (blue) or NLS-WT Tgm2 (orange) treated stage 13 (non-migratory) embryos were grafted into wild-type stage 13 host embryos. **b,** Embryos displaying the results of the grafts; vertical white lines indicate migrated distance; scale bar, 150 μm; conditions as indicated. **c,** Percentage of embryos displaying neural crest migration. Histograms show mean and standard deviation, two-tailed t test with Welch’s correction, \*\**P* =0,0015, n_(NLS-GFP)_ = 9 embryos; n_(NLS-WT Tgm2)_ = 19 embryos. **d,** Schematic of extrinsic stress AFM experiments, images of embryos being compressed from non-migratory (stage 13) to pre-migratory stages (stage 15) with a 90-μm bead attached to an AFM cantilever (bead, black circle) are shown; scale bar, 250 μm; cleared and flat-mounted imaging scheme is also shown (details in Methods). **e,** Tgm2-GFP (no NLS-tagged) (orange) in control or extrinsically stressed embryos; nuclei are shown in cyan; scale bar, 20 μm. **f,** Immunostaining for histone serotonylation (magenta) in control or extrinsically stressed embryos; scale bar, 20 μm; **d-f**, conditions as indicated. **g,** Nuclear/cytosolic ratio of Tgm2-GFP intensity; red lines show mean, error bars represent standard deviation; two-tailed t test with Welch’s correction, \*\*\**P* < 0.0006. **h,** Normalised histone serotonylation fluorescence intensity; spread of data from the indicated conditions is shown, red lines represent median and whiskers represent interquartile ranges, two-tailed Mann Whitney test, \*\**P* = 0,0019, n_(control)_= 9 embryos; *n*_(stressed)_ = 9 embryos. **i,** *In situ* hybridization, lateral view of *sox8*-hybridized embryos; scale bar, 150 μm; conditions as indicated. **j,** Percentage of embryos displaying neural crest migration, histograms represent mean and error bars standard deviation, one-way ANOVA, \*\*\*\**P* =0,0001, at least 9 embryos per condition were used. Panels in b, e, f and i are representative examples of at least three independent experiments. CI = 95%.

We next sought to explore the nature of the transcriptional program that is mediated by histone serotonylation in response to substrate stiffening. For this, we performed Chromatin Immuno Precipitation (ChIP) against endogenous histone serotonylation from freshly dissected pre-migratory neural crests (**Fig. 5a**; **Methods**). Following sequencing, a list of candidates predicted to be regulated by histone serotonylation (enriched within proximal promoters) was obtained (**Fig. 5b**). This is in agreement with the previously described permissive transcriptional activities of this mark^4,12,22^. Motif analyses identified consensus binding sites for families of transcription factors that are associated with histone serotonylation^6^, as well as others required for neural crest epithelial to mesenchymal transition (EMT), one of the first steps of their migration (i.e., Hif1, Twist, Zeb)^23,24^ (**Fig. 5c**). We next validated whether the transcription of these candidates relies on histone serotonylation. For this, we performed RNA-seq experiments from isolated neural crest cells obtained from embryos treated with wildtype or mutant histone H3 (H3Q5A) (**Supplementary Fig. 7a**; **Methods**). Several transcripts were found to either increase or decrease (**Supplementary Fig. 7b**), but owing the permissive roles of histone serotonylation, we focused on the list of downregulated candidates. By comparing this list with the one obtained from our ChIP-seq experiments, we identified 91 genes whose transcription relies on the permissive activities of histone serotonylation (**Fig 5d**). To further filter this list, we next dissected what portion of the identified transcripts are regulated by histone serotonylation in response to mesoderm stiffening. We performed RNA-seq from isolated neural crest cells obtained from wildtype embryos or embryos treated with ca-Mypt1 to soften their mesoderm (**Supplementary Fig. 7c**). We obtained several candidates whose expression was up or down-regulated upon substrate softening *in vivo* (**Supplementary Fig. 7d**). However, as discussed above, owing to the permissive roles of histone serotonylation, we focused our analysis only on the list of downregulated transcripts. Global comparisons of this list with the one obtained from our alignments of histone serotonylation ChIP-seq and RNA-seq datasets revealed a refined list of 37 common candidates whose transcription could be considered to be directly regulated by histone serotonylation in response to substrate stiffening (**Fig. 5e, f**). Gene ontology analysis indicated that 29 of these candidates are related to cell migration (**Fig. 5g**) and that 14 of them are related to EMT, an essential step for neural crest cell migration (**Fig. 5g**). At least 10 of these 29 candidates (*epb41l5.S*^25–27^, *ephb4.S*^28,29^, *erbb2.L*^30^, *gpc1.L*^31,32^, *lrp8.L*^33^, *myh9.S*^34,35^, *rasa3.S*^36^, *smad1.S*^37^, *wnt9a.S*^38^, and *yme1l1.S*^39,40^) are known to be related to neural crest migration. Furthermore, our study also revealed candidates whose expression in migrating neural crest has been reported, however, their roles in migration have not yet been elucidated. In summary, these data revealed a cell migration specific transcriptional module that is activated at the onset of neural crest migration via mechanical regulation of histone serotonylation without affecting major neural crest cell fate markers **(Fig. 5e, h).** Further validation of these candidates in the neural crest and other tissues will assist in providing a more complete understanding about the mechanical modulation of histone serotonylation during CCM.

**Figure 5.**
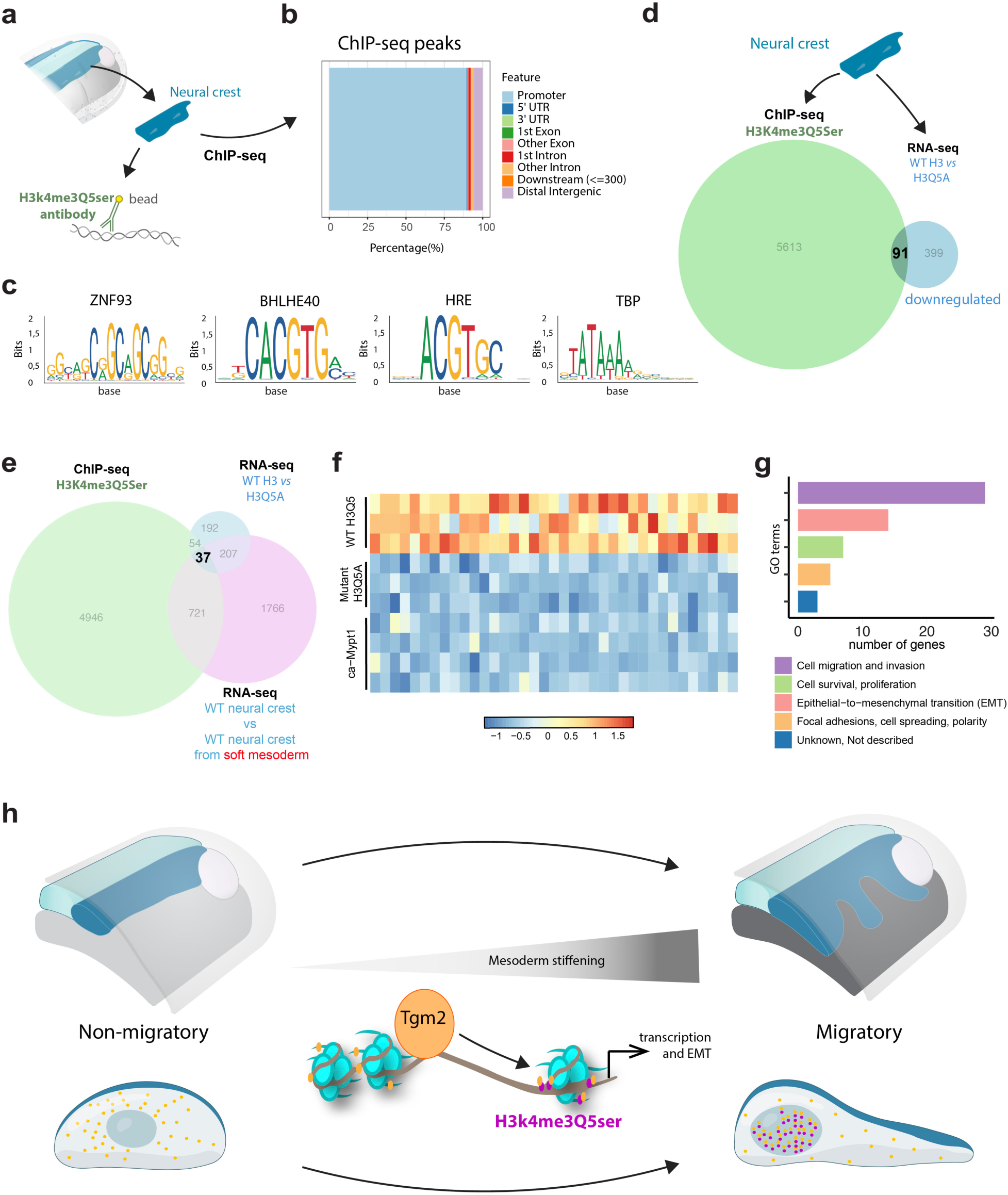
Description of transcriptional modulation by histone serotonylation in response to mesoderm stiffening. **a**, Schematic showing the workflow of ChIP-seq against endogenous H3k4me3Q5ser from isolated neural crest cells (peak calling using q-value cutoff of 0.05). **b,** Chart displaying the distribution of H3k4me3Q5ser ChIP-seq peaks across genomic regions. **c,** Motif-enrichment analysis of significantly altered sequences obtained from ChIP-seq analysis highlighting consensus binding sites. **d,** Schematic showing the workflows of ChIP-seq and RNA-seq analyses from isolated neural crest cells. Genes identified through ChIP-seq peak annotation were compared with downregulated genes from RNA-seq differential expression analysis obtained after comparing wild-type and mutant histone H3 (H3Q5A), 91 potential target genes were identified [Log2FC < 0.58 and FDR-adjusted p-value < 0.05]. **e,** Comparison of the target lists from **d** with downregulated genes from RNA-seq differential expression analysis of wild-type neural crests from CA-MYPT treated-embryos, 37 candidates were identified [Log2FC < 0.58 and FDR-adjusted p-value < 0.05]. **f,** Heatmap visualization of the differential expression of 37 candidate genes regulated by histone serotonylation in response to mesoderm mechanics, using z-scores to indicate standardized expression levels across the analysed conditions. **g,** GO-term enrichment of the 37 candidates. **a-g**, all experiments were repeated at least two times. **h,** Model showing that mesoderm stiffening promotes nuclear shuttling of Tgm2 and histone serotonylation to allow chromatin fine-tuning and with that the activation of a transcriptional program which is essential to trigger neural crest CCM.

## Outlook

The mechanical influence of gene transcription via nuclear shuttling of transcription factors (TFs), such as Yap1/Taz, has previously been proposed^41^. Yet, whether and how the mechanical microenvironment modifies the chromatin landscape to influence gene expression is just beginning to be studied in detail^19,42–44^. As a consequence, very little remains known regarding the mechanical regulation of gene transcription within physiologically relevant contexts (*in vivo*)^19,45,46^. Our results here provide *in vivo* evidence demonstrating a previously uncharacterized mechanism in which substrate mechanics promote chromatin modifications and, with that, support a defined transcriptional program for the initiation of CCM. Our combination of *in vivo* and *ex vivo* experiments, together with mechanical and molecular perturbations, show that serotonylation of histone H3 occurs within the cranial neural crest, enabling the onset of its collective migration *in vivo*. Furthermore, we found that histone serotonylation is mediated by mechanical stiffening of the mesoderm, a tissue that acts as the migratory substrate for neural crest cells *in vivo*. Mechanistically, we found that mesoderm stiffening promotes nuclear shuttling of Tgm2, the primary *writer* of histone serotonylation.

Histone serotonylation has only recently been discovered, and very little remains known regarding its physiological functions^4,47,48^. Therefore, its participation as a molecular mediator of the transcriptional response of migratory cells to substrate stiffening was not addressed previously. Similarly, whether and how environmental cues modulate histone serotonylation levels/genomic enrichment have remained elusive. Hence, our observations regarding the mechanical regulation of histone serotonylation in the neural crest by substrate stiffening appear to be of great relevance. Likewise, the nuclear shuttling of Tgm2 was published previously in 1983^49^, yet while a large body of evidence placed it as a modifier of histones^50–52^, its role in mediating histone serotonylation was only recently described. Mechanical control of Tgm2 nuclear shuttling has not been reported previously and its role in neural crest CCM has not been addressed to date. Since mechanical modulation of chromatin state is only beginning to be studied in physiological contexts, our work also contributes with mechanistic insights to this growing field of “mechanoepigenetics.”

In addition, we found that histone serotonylation directly regulates the expression of a defined subset of targets that support the onset of collective cell migration via EMT. It was previously shown that substrate stiffening synchronizes morphogenesis by triggering the onset of neural crest CCM *in vivo* via EMT^5^. However, whether this occurs via direct regulation of chromatin, and a detailing on the transcriptional modules regulated by mesoderm stiffening, remained elusive. Further validation of the candidates identified here will shed light on the mechanical control of EMT *in vivo*. In addition, it was previously shown that histone serotonylation reinforces, but does not fully modify H3K4me3-related transcriptional programs. Our screening agrees with this idea, as we identified a rather low number of candidate genes that are directly targeted by histone serotonylation in response to mechanical stimuli. These observations suggest that, to some extent, histone serotonylation fine-tunes the neural crest transcriptional program to switch from a non-migratory to a migratory cell state without affecting its original commitment to a neural crest fate.

Tissue mechanics, EMT, and collective cell migration are relevant for embryonic development, tissue regeneration and cancer metastasis^5,53–56^, and there is a growing interest in the potential roles of histone serotonylation in physiology and disease^47,57–59^. Therefore, we believe that our results open up interesting avenues of research in which roles for mechanical regulation of histone serotonylation can be explored in other biological contexts.

## Acknowledgments

We thank Dr Ana Patrícia Ramos (Pol, TUD) and Pablo Strobl-Mazulla (INTECH, Argentina) for helpful comments to the manuscript. The authors also acknowledge Gulbenkian’s and the Physics of Life Advanced imaging, Genomics, Bioinformatics and Aquatic animal facilities. Work at E.H.B. lab was funded by the European Research Council (ERC) under the European Union’s Horizon 2020 research and innovation programme (grant agreement No. 950254), EMBO Installation Grant (4765), la Caixa Junior Leader Incoming (94978), and EMBO-YIP (5248). E.H.B. also acknowledge the support from Fundação Calouste Gulbenkian (start-up fund I-411133.01) and the Deutsche Forschungsgemeinschaft (DFG, German Research Foundation) under Germany’s Excellence Strategy (EXC 2068, 390729961, Cluster of Excellence Physics of Life of TU Dresden). J.E.S. was supported by an FCT doctoral fellowship (UI / BD / 152251 / 2021) and I.M. work was supported by NIH R01MH116900 and Funds from the Howard Hughes Medical Institute.

## Author contributions

E.H.B. conceived the project. J.E.S. performed most of the experiments and data analyses with help from E.H.B., M.R., and J.A.E. X.W., I.M. and A.G.K. performed bioinformatic analysis. I.M. generated plasmids and designed ChIP protocols. J.E.S. and E.H.B wrote the manuscript and prepared the figures. All the authors edited the manuscript. E.H.B supervised the project and secured funds.

## Data and materials availability

All data supporting our findings are provided in the main text or supplementary material. Any other relevant data are available from the corresponding author upon reasonable request.

## MATERIALS AND METHODS

All animal experiments and euthanasia were reviewed and approved by the Ethics Committee and Animal Welfare Body (ORBEA) of the Instituto Gulbenkian de Ciência (IGC) and complied with the Portuguese (Decreto-Lei n° 113/2013) and European (Directive 2010/63/EU) legislation.

### *Xenopus laevis* manipulation and embryo generation

Adult frogs were maintained in a temperature and light-controlled environment (18 °C and 12h light/12h obscurity). *Xenopus laevis* embryos were obtained as previously described^23^. Briefly, ovulation of wild type mature females was induced by injecting human chorionic gonadotropin (MSD Animal Health, Chorulon). Collected eggs were fertilized *in vitro* by mixing with a sperm suspension in 0.1x Marc’s modified Ringer medium (MMR: 10 mM NaCl, 0.2 mM CaCl_2_·2H_2_O, 0.2 mM KCl, 0.1 mM MgCl_2_·6H_2_O, and 0.5 mM HEPES, adjusted to pH 7.1–7.2). Testes were dissected from mature wild type males. Embryos were incubated between 12 and 23°C in 0.1x MMR until reaching the desired stages, according to established developmental tables^60^.

### *In situ* hybridization and *in vitro* probe transcription

Embryos were fixed and processed following previously established protocols^61^. Digoxigenin-labelled antisense probe against *sox8*^62^ was transcribed with a Riboprobe® *in vitro* Transcription System (Promega, P1420), according to the manufacturer’s instructions. DIG RNA Labeling mix (Roche 11277073910) was used. RNA cleanup was performed via a RNeasy kit (Qiagen) and the RNA was eluted in RNase-free H_2_O (Ambion). To obtain the t*gm2* probe, a PCR fragment of ∼750 bp was amplified from the Xenopus *tgm2* full length by using primers containing the T7 promoter for easy transcription. Primers 11-12 (Supplementary Table 1) for the antisense probe; and primers 13-14 (Supplementary Table 1) for the sense probe were used. Annealing temperature for PCRs was 62°C. Transcription was performed using the Promega Riboprobe® *in vitro* transcription system (P1440 for the T7 RNA polymerase kit) as aforementioned. Probes were used at a concentration of 3 μg/ml in hybridization buffer^61^. The signal was revealed using the alkaline phosphatase detection reagents NBT/BCIP at a proportion of 4.5/3.5 μl/ml (Roche, 11383221001).

### Morpholino and mRNA injections and reagents

Eggs were de-jellied in a 2% cysteine solution in 0.1x MMR (w/v) with 500 μl of 5 M NaOH and transferred to ficoll 5% in 0.45× MMR (Sigma-Aldrich, P7798). Morpholino and mRNA injections were performed by targeted microinjections using glass needles calibrated to inject 10nL when using 20psi of pressure in a 0.2 seconds gas pulse. A cell microinjector (MDI, PM1000) was used. For neural crest-targeted injections, 8-cell stage embryos were injected near the division point of a dorsal and a ventral blastomeres of the animal pole. When required, cells were fluorescently tagged with mRNA of nuclear RFP, membrane RFP and/or membrane GFP (at 250 pg per blastomere per construct); these transcripts were generated *in vitro* by using the mMESSAGE mMACHINE SP6 kit (Thermo-Fisher, AM1340), according to manufacturer’s instructions. Tgm2-MO was synthesized by GeneTools (sequence in Supplementary Table 1) and injected at 7 ng/blastomere. WT H3, H3Q5A, NLS-WT Tgm2 and NLS-mutant Tgm2 were previously validated^4^. Tgm2 ORF2 – XORFeome v1.0 (Xenopusresource) was used to generate a Tgm2 (full-length) insert. All PCR products were sub-cloned into a pCS2+8CeGFP (Addgene, 34952) plasmid, transcribed as described above. PCR templates were generated by using the primers 1-2 (Supplementary Table 1) for Tgm2 (full-length); primers 3-4 (Supplementary Table 1) for WT H3; primers 5-6 (Supplementary Table 1) for H3Q5A; primers 7-8 (Supplementary Table 1) for NLS-WT Tgm2; primers 9-10 (Supplementary Table 1) for NLS-mutant Tgm2. Injections were as follows: H3 constructs were injected at 750 pg per blastomere and Tgm2 constructs at 1ng per blastomere; Note that these concentrations were decided as follow. WT H3 or NLS-WT Tgm2 constructs were optimised up to a concentration until we did not detect any differences on the levels of histone serotonylation between those treatments and the WT-uninjected neural crests; then the mutant constructs (H3Q5A or NLS-mutant Tgm2) were injected at the same concentrations. Tgm2-GFP (full-length) was injected at 100 pg per blastomere; Vinculin- cTERwas injected at 250 pg per blastomere. For targeted mesoderm injections, 1 ng of CA-MYPT ^20^ was injected into dorso-vegetal blastomeres at the 16-cell stage.

### Neural crest dissection

Cephalic neural crest cells were dissected from embryos at stages 16 and 17 and cultured *ex vivo* as previously described^61^. Briefly, devitellinized embryos were placed in a dish containing plasticine and filled with 0.3x MMR. Embryos were immobilized by securing them with plasticine. A hair knife was used to remove epidermis and explant the underlying neural crest into dishes containing Danilchik’s for Amy 1x medium (DFA: 53 mM NaCl, 5 mM Na_2_CO_3_, 4.5 mM potassium gluconate, 32 mM sodium gluconate, 1 mM MgSO_4_·7H_2_O, 1 mM CaCl_2_·2H_2_O, 0.1% BSA (w/v), adjusted to pH 8.3 with 1 M Bicine). Clusters adhered to glass-bottom dishes coated with 62.5 μg ml^−1^ of fibronectin (Sigma-Aldrich, F1141) for 30-60 min.

### Dispersion assay

A spreading assay was used to analyse the migration of neural crest *ex vivo.* Explants were allowed to attach and their migration and dispersion was recorded by time-lapse microscopy. These assays have been extensively used as a readout of EMT and allow the analysis and comparison of several motility parameters among conditions^61^.

### Graft experiments

Wild-type or injected neural crests were dissected as aforementioned. Then the donor neural crest was carefully placed on a host embryo by using a hair knife and held in place by a small piece of coverglass (∼1×1 mm). After approximately 1h, the coverslip was removed. The embryos were cultured in 0.3x MMR and fixed when reaching the stage required for each experiment.

### Polyacrylamide (PAA) hydrogels preparation

Soft gel mixes contained: 550 μl of 7.6 mM hydrochloric acid (HCL), 350 μl of double-distilled water (ddH_2_O), 0.5 μl N,N,N′,N′-tetramethylethylenediamine (TEMED) (Sigma), 20 μl 2% bis-acrylamide (BioRad), 70 μl of 40% acrylamide (BioRad), 20 μl 0.1 M NHS (N-hydroxysuccinimide, Sigma-Aldrich), 5 μl of 200 nm diameter beads resuspended at 0.2 μM (Invitrogen) and 5 μl of 10% ammonium persulfate (GE HealthCare) (freshly prepared). Stiff gels mixes contained: 550 μl of 7.6 mM HCL, 282 μl of ddH_2_O, 0.5 μl of TEMED (Sigma), 25 μl 2% bis-acrylamide (BioRad), 137 μl of 40% acrylamide (BioRad), 20 μl of 0.1 M NHS (N-hydroxysuccinimide, Sigma-Aldrich), 5 μl of 200 nm diameter beads resuspended at 0.2 μM (Invitrogen) and 5 μl of 10% ammonium persulfate (GE HealthCare) (freshly prepared). 13-mm diameter × 0.1 mm hydrophobic glass coverslips were coated for 15 min at room temperature with PlusONE Repel-Silene ES (GE Healthcare), carefully air-dried with an air pistol and used immediately. A 12-μl drop of PAA mix was added on a hydrophilic glass dish (FD5040-100) and immediately covered with pre-coated hydrophobic coverslips to promote the attachment of gels to glass slides. PAA polymerization proceeded for 45 min at room temperature in a humidifier chamber. The coverslip was carefully removed, and gels were washed three times for 2 min with PBS 1x.

### Gel functionalization

Fibronectin was covalently linked to the hydrogels with 0.2 M EDC ((1-ethyl-3-(3-dimethylaminopropyl)carbodiimide hydrochloride), Calbiochem), 0.1 M NHS (N-hydroxysuccinimide, Sigma-Aldrich), in 0.1 M MES buffer (in milliQ water, pH 5.0, 2-(N-morpholino)-ethane sulfonic acid, Sigma-Aldrich). After washing twice with PBS, gels were incubated in a humidifier with 10 mg/ml fibronectin for 1h45 min at room temperature. Fibronectin was washed with PBS and the crosslinking-reaction was quenched by incubating the gels for 15 min with 0.32% ethanolamine (Sigma-Aldrich) in PBS. Fluorescent Fibronectin (Cytoskeleton, Inc., HiLyte 488) was used to determine gel functionalization. Neural crest cells were dissected and placed on the prepared gels immersed in DFA medium for 2-3 hours until they attached and spreaded on gels.

### Cryosectioning

Embryos were fixed at the desired stages in a solution containing 4% formaldehyde and 0.3% TritonX-100 in 1X PBS. Embryos were fixed for 2-3h at room temperature, followed by two 5min washes in phosphate buffer (0.2 M NaH_2_PO_4_·H_2_O and 0.2 M K_2_HPO_4_, pH 7.4). Then, samples were incubated for 2h at room temperature in 15% sucrose (wt/vol; Sigma-Aldrich) in phosphate buffer and for 1h at 42 °C in a 8% gelatin (Sigma-Aldrich) and 15% sucrose solution in phosphate buffer (w/v). Embryos were embedded and oriented in gelatin solution and gelatin blocks containing the embryos were snap frozen at −80 °C with pre-chilled isopentane. Samples were sectioned in 30-μm slices using a cryostat (CM-3050S, Leica) and collected in SuperFrozen Slides (VWR International). The slides were dried for at least 6h and the gelatin was removed by two 10min washes with 1x PBS at 42 °C. Slides were processed for immunostaining, as described below.

### Immunostaining in glass, hydrogels, cryosections and flat-mounted embryos

Fibronectin (mAb 4H2 anti-FN, DSHB)^63^, H3k4me3Q5ser (Merck-Millipore ABE2580)^4^, H3k4me3 (Cell signalling 9751)^64^ and Tgm2 (Abcam ab2386)^4^ primary antibodies were used for immunostaining of cryosections, flat-mounted embryos and/or *ex vivo* neural crest explants. For H3k4me3Q5ser, H3k4me3 and Tgm2 detection *ex vivo,* explants were permeabilized with PBS 0.1% Triton X-100 for 5 min at room temperature. Blocking was performed with 10% normal goat serum (NGS) for 30 min at room temperature. Cryosections were blocked for 1 hour with 10% NGS. Antibodies were diluted in blocking solution. Anti-fibronectin was used at 1:40 for cryosections and flat-mounted embryos; Anti-H3k4me3Q5ser was used 1:1000 for cryosections, 1:400 for explants, and 1:100 for flat-mounted embryos; Anti-H3k4me3 was used 1:200 for cryosections and 1:400 for explants; Anti-Tgm2 was used 1:100 for explants. All primary antibodies were incubated overnight at 4 °C and washed three times with 0.1% PBS-T (PBS, 0.1% Tween-20). Alexa-fluor secondary antibodies used in this study were anti-rabbit 488 (LifeTechnologies #A11034), anti-rabbit 555 (LifeTechnologies (#A21429), anti-rabbit 647 (LifeTechnologies #A31573), anti-mouse 555 (LifeTechnologies #A31570) and anti-mouse 647 (LifeTechnologies #A32728). Secondary antibodies were diluted 1:350 in blocking solution with 1/700 DAPI (for nuclear staining). The samples were incubated in this mix for 2-3h at room temperature and washed three times with 0.1% PBS-T. Immunostaining on hydrogels proceeded as described above but washes were replaced by gentle rinses. MOWIOL (EMD Millipore) was used as the mounting medium. For immunostaining in flat-mounted embryos, whole embryos were fixed using 4% formaldehyde in 0.3% PBS-T (PBS, 0.3% Triton X-100) for 3 hours. Dorsal halves of fixed embryos were dissected using a 15°, 5-mm depth microsurgical knife (MSP 7516). Dissected halves were washed with PBS and blocked for 1 hour with 10% NGS. Anti-fibronectin and anti-H3k4me3Q5ser antibodies were incubated overnight at 4°C, followed by three washes with 0.3% PBS-T. Secondary antibodies were incubated for 3 hours and excess antibody was removed by washing three times with 0.3% PBS-T. Nuclei were stained with DAPI in a mixture with the secondary antibodies. Finally, embryos were fixed with 4% formaldehyde in 0.3% PBS-T for 30 min, washed overnight in methanol and flat-mounted for imaging in clearing mix (two volumes benzyl benzoate and one volume benzyl alcohol; Sigma-Aldrich). To detect fibronectin on gels, functionalized gels were washed twice with PBS and incubated for 1 hour at room temperature with anti-human fibronectin (Sigma S3648) diluted 1:500 in PBS. Gels were washed three times with PBS and incubated for 1 hour at room temperature with 1:350 Alexa-fluor anti-Rabbit 488. Gels were washed three times with PBS and maintained in 1% formaldehyde in PBS.

Alexa-fluor (Thermo-Fisher) secondary antibodies were diluted 1:350 in blocking solution with 1/700 DAPI (for nuclear staining). The samples were incubated in this mix for 2-3h at room temperature and washed three times with 0.1% PBS-T. Immunostaining on hydrogels proceeded as described above but washes were replaced by gentle rinses. MOWIOL (EMD Millipore) was used as the mounting medium. For immunostaining in flat-mounted embryos, whole embryos were fixed using 4% formaldehyde in 0.3% PBS-T (PBS, 0.3% Triton X-100) for 3 hours. Dorsal halves of fixed embryos were dissected using a 15°, 5-mm depth microsurgical knife (MSP 7516). Dissected halves were washed with PBS and blocked for 1 hour with 10% NGS. Anti-fibronectin and anti-H3k4me3Q5ser antibodies were incubated overnight at 4 °C, followed by three washes with 0.3% PBS-T. Secondary antibodies were incubated for 3 hours and excess antibody was removed by washing three times with 0.3% PBS-T. Nuclei were stained with DAPI in a mixture with the secondary antibodies. Finally, embryos were fixed with 4% formaldehyde in 0.3% PBS-T for 30 min, washed overnight in methanol and flat-mounted for imaging in clearing mix (two volumes benzyl benzoate and one volume benzyl alcohol; Sigma-Aldrich). To detect fibronectin on gels, functionalized gels were washed twice with PBS and incubated for 1 hour at room temperature with anti-human fibronectin (Sigma S3648) diluted 1:500 in PBS. Gels were washed three times with PBS and incubated for 1 hour at room temperature with 1:350 Alexa-fluor anti-Rabbit 488. Gels were washed three times with PBS and maintained in 1% formaldehyde in PBS.

### Microscopy and time-lapse live imaging

#### Time-lapse imaging

Images for dispersion assays were acquired every 5 min at 18 °C with a Leica Stellaris 5 confocal system, using a 20x objective. Camera (DFL420, Leica), filter wheels and shutters were controlled by built-in Leica software.

#### In situ hybridization and grafts imaging

*In situ* hybridized and grafted embryos were mounted on an agarose dish with small depressions to facilitate acquisition and imaged with an USB Dino-Eye eyepiece camera (Dino-Lite, AM7025X), mounted onto a stereoscope with cold light at 2.5× magnification, using DinoCapture v2.0 (Dino-Lite); Some *in situ* hybridized embryos were imaged using Zeiss SteREO Lumar equipped with a Hamamatsu Orca-Flash 2.8 camera, controlled with the MicroManager v1.14 software. All images were captured at room temperature.

#### Immunofluorescence imaging

Most of the images were acquired at room temperature using a Zeiss LSM900 or LSM980 system, equipped with two PMT and one GaAsPand, a 40x W objective (C-Apochromat 40x/1.1 W Corr M27, Zeiss); Camera, filter wheels and shutters were controlled by Zeiss’s ZEN Blue v.3.0. Some images were acquired with a Leica Stellaris 5 confocal system, using 20x or 40x objectives. Camera, filter wheels and shutters were controlled by built-in Leica software.

#### *In vivo* AFM measurements

All AFM measurements were performed as previously described^5,21,65^ by using a FLEX-ANA (Nanosurf) automated AFM device, fitted with a x–y motorized stage and an automated software for experimental setup and analysis (ANA Software, Nanosurf). Cantilevers were coated with roughly 10 μm diameter colloidal spheres (CP-qp-SCONT-BSG-B-5, sQube). Cantilevers were mounted on the AFM device and their spring constants were calculated using the thermal noise method^66^. Only cantilevers with spring constants between 0.01 and 0.03 N m^−1^ were selected. Embryos were mounted in a plasticine dish and indentations were performed with the following modulation parameters: maximum indentation force, 10 nN; approach speed, 5 μm s^−1^; retraction speed was 55 μm s^−1^ and sample rate, 2,400 Hz. Extrinsic stress experiments were performed as previously described^5^ with a Nanowizard IV (Brucker/JPK) atomic force microscope. Briefly, stress was applied to the migratory path of the neural crest with a 0.1 N/m cantilever that was coated with a bead of 87.3 ± 1.1 μm. A sustained force of 80 nN was applied from stage 13 (non-migratory) until reaching early pre-migratory stages (stage 15). Contralateral side were used as controls. An image was acquired before and after each indentation, to generate displacement maps in the ImageJ iterative PIV basic plugin. After compression, embryos were rapidly fixed for *in situ* hybridization against *sox8*. Scan head, stage and camera, are controlled by the JPK software (8.0.144).

### Data analysis and image treatment

#### AFM data

Force–distance curves were analysed with a Hertz model for a spherical indenter as previously described^21,65^, for defined indentation depths of 3 μm, which has been shown to measure the mechanical properties of developing tissues^67–70^. Measurements for each experiment were pooled and statistically analysed. Scale bars were calculated with ImageJ and applied with Adobe Illustrator.

#### Immunofluorescence data

For each explant, the fluorescence intensity of 5 cells were measured, normalized to the nuclear marker, subjected to background subtraction, averaged and min-max normalized in Fiji (ImageJ 2.1.0). Z-stacks and maximum projections were created using ImageJ software. Addition of scale bars and pseudocolour were applied with ImageJ and/or Adobe Photoshop. *<u>In situ</u>* <u>data.</u> In Fig. 1g, 2a, 2g, 5i and Supplementary Fig. 6e, the background was pseudocoloured in Photoshop and addition of scale bars were applied with ImageJ and/or Adobe Photoshop.

### *In vivo* analysis of neural crest migration

For *in situ* hybridization and grafted embryos, the displacement of neural crest cells was obtained by normalizing the length of the migratory stream against the total dorso-ventral length of the embryo. Length values were obtained in Fiji (ImageJ 2.1.0) using the measurement tool and further analysed as described in Statistical analysis.

### *Ex vivo* analysis of neural crest spreading

For dispersion assays, an ImageJ-based custom-made Delaunay-triangulation plugin^71^ was used to calculate the distance between neighbouring cells (available upon request). Data were further analysed as described below in Statistical analysis.

### Image treatment

Substacking, z-projections (maximum intensity), rotations, time colour-coded projections, were performed in Fiji (ImageJ 2.1.0). General treatment, including adjustment of contrast and brightness, resizing, pseudocolouring, addition of scales and overlay of text in images were performed in Fiji and/or Adobe Photoshop CS6. In projections on cryosections, the background with neighbouring tissues was coloured in grey for clarity purposes and without interfering with the sample itself, using Fiji.

### sqRT-PCR

Neural crest explants from pre-migratory stages (stg 17–18) were collected and processed for RNA extraction using the RNeasy Mini kit (Qiagen, 74104). Briefly, 5 ng of isolated total RNA were used as a template for the RT performed using SuperScript IV (Invitrogen,18090050), according to manufacturer’s instructions. To run the PCRs, 2 ng of resultant cDNA in 25 μl of reaction were used for the genes of interest, using Q5 Hot Start High-Fidelity DNA Polymerase (NEB, 174M0493S) with the following primers: for *tgm2* primers 15-16 (Supplementary Table 1); for *ef1α* primers 17-18 (Supplementary Table 1). PCR conditions were as follows: (1) 98 °C for 30 s (initial denaturation); (2) 98 °C for 10 s (denaturation); (3) 30 s (annealing) at 64 °C (*ef1α*), 67 °C (*tgm2*); (4) 72 °C for 20 s (32× cycles from step 2 to 4); and, finally, 72 °C for 55 s for final extension. The amplified fragments were resolved in agarose 0.8% (w/v) gel electrophoresis together with 1 Kb Plus Ladder (Invitrogen, 10787026). While *tgm2* primers were originally designed, *ef1α* primers were previously validated^72^.

### RNA-seq procedures and analysis

The quality of the extracted RNA was assessed using HS RNA Screen Tape Analysis (Agilent Technologies). Libraries were generated by SMART-Seq2 and a Fragment Analyzer Advanced Analytical Technologies (AATI) was used for their quantification and quality control. Sequencing was performed in a NextSeq500 Sequencer (Illumina) using 100 SE high throughput kit. Sequence information was extracted in FastQ format using the bcl2fastq v2.19.1.403 (Illumina). Raw sequencing fastq files were assessed for quality and adapter content with FastQC v0.12.1 and multiqc v1.14. Adapters were trimmed and reads with low-quality or length less than 10 bases were discarded using cutadapt v4.4^73^ (with the following parameters: -b CTGTCTCTTATACACATCT -b GTATCAACGCAGAGTACT -b TTTTTTTTTTTTTTTTTTTTTTTTTTTT -b AAAAAAAAAAAAAAAAAAAAA -m 10 -q 20). The rRNA content were filtered further with SortMeRNA v4.3.6^74^ using as reference the eukaryotic 18S and 28S rRNA sequences. The trimmed, rRNA depleted samples were aligned to the reference genome X. laevis v10.1 using hisat2 v2.2.1 with default parameters^75^. Subsequently, the resulted SAM files were converted into BAM files and sorted using samtools v1.5. Transcript abundance was quantified using featureCounts v2.0.6^76^, generating a raw count matrix of mapped reads per gene. Raw counts were normalized to account for technical variations, including library size, using the “DESeq2” package in R^77^. Differential gene expression analysis identified genes with statistically significant expression changes between conditions, applying thresholds of |Log2FC| > 0.58 and FDR-adjusted p-value < 0.05. Volcano plots were generated to visualize the differential gene expression using the “ggplot2” package in R. Additionally, raw counts for each condition were transformed into z-scores for downstream visualization purposes.

### ChIP-seq procedures and analysis

ChIP was performed according to standard procedures established for *X. laevis* embryos^78,79^. Briefly, stage 17 wild type neural crests were dissected from 100-140 *X. laevis* embryos, fixed for 30 min at room temperature and quenched. Samples were washed thoroughly before being subjected to lysis and were sonicated for 40 min at 30% power (30 s On 30 s Off) in a QSonica Q800R3 sonicator. Then, samples were incubated with 5 μg of Anti-H3k4me3Q5ser antibody (Merck-Millipore ABE2580) bound to M-280 Dynabeads (Invitrogen) on a rotator at 4 °C overnight to precipitate a fraction of the total sonicated extract. The following day, immunoprecipitates were washed, eluted and reverse-crosslinked. Following RNA and protein digestion, DNA fragments were purified using a Qiagen PCR purification kit. Sequencing libraries were generated by using NEBNext Ultra II DNA Library Prep Kit (Illumina). The quality of the libraries was assessed using a High Sensitivity DNA assay of Fragment Analyzer (Agilent Technologies). Sequencing was performed in a NextSeq2000 Sequencer (Illumina) using 100 cycles, 50M reads of sequencing depth, 50 bp PE high throughput kit.

Raw sequencing reads were first subjected to quality control using FastQC v0.12.1 and multiqc v1.14 to ensure data integrity. Adapter sequences were removed with cutadapt 4.4^73^, using the following 3′ adapter sequences: AGATCGGAAGAGCACACGTCTGAACTCCAGTCA for Read 1 and AGATCGGAAGAGCGTCGTGTAGGGAAAGAGTGT for Read 2. The trimmed reads were aligned to the reference genome X. laevis v10.1 using Bowtie2 v2.4.4^80^ with the following parameters: --fr --end-to-end --very-sensitive --no-discordant --no-mixed. SAM files were converted into BAM files and sorted using samtools v1.5. Further filtering to remove duplicates, multimappers, and unmapped reads was performed using sambamba v1.0.0 with the following command: sambamba ^81^ view -h -t 2 -f bam -F "[XS] == null and not unmapped and not duplicate". Finally, the filtered BAM files were used for peak calling. Peaks were called for individual replicates of the IP and input samples with MACS2 v2.2.8^82^ using a q-value cutoff of 0.05. The peak files were filtered by intersecting the peaks called in the corresponding input sample using bedtools v2.26.0^83^ with the following command: bedtools intersect -v -f 0.3 -r.

The peaks were associated with gene promoter regions using bedtools intersect (-nonamecheck -wo), reporting overlapping peaks with manually extracted transcription start site (TSS) regions of the custom genome. The fasta sequences of the selected peaks were extracted using the bedtools getfasta command. Motif enrichment analysis was conducted using the Simple Enrichment Analysis (SEA) tool from the MEME suite^84^, with the JASPAR2022 CORE vertebrates non-redundant v2 database as the reference for motif discovery.

### Comparison of RNA-seq and ChIP-seq datasets

The gene list obtained from ChIP-Seq peak annotation was compared with the set of downregulated genes identified in the RNA-Seq differential expression analysis in different conditions. The overlap between these gene sets was visualized using the package “eulerr” in R. The z-scores of the intersecting genes were displayed in a heatmap generated with the “pheatmap” package. For functional enrichment analysis, the final gene set was annotated with GO terms and a bar plot illustrating the distribution of genes across predefined categories was generated using the “ggplot2” package.

### Statistical analysis

The experimenters were not blinded in data acquisition, treatment, or statistical analysis due to the nature of the experimental procedures, since only well-injected, non-contaminated and viable embryos/cell clusters were considered for analysis. For any of the mentioned cases, after selections, all parameters were measured at random. Data outliers were removed using the statistical Grubbs’ test and/or ROUT method computed using GraphPad Prism 6.0. Majority of datasets presented no statistical outliers. The appropriate inferential statistics were computed using GraphPad Prism and are annotated in the figure legends. Each experiment was repeated at least three times, using embryo batches from different females. Every set of data was tested for normality test using the, d’Agostino–Pearson and/or Shapiro–Wilk test in GraphPad Prism. For paired comparisons, significances were calculated in GraphPad Prism with a Student’s *t*-test (two-tailed, unequal variances) when the distributions proved to be normal. If a data set did not pass the normality tests, the significances were calculated with Mann–Whitney (two-tailed, unequal variances). For multiple comparison of data with normal distribution unpaired one-way analysis of variance (ANOVA) with Bonferroni test correction was performed, while non-normal distribution data sets were analysed with Kruskal–Wallis corrected with Dunn’s test. Differences were significant when the two-tailed *p* value was ≤ 0.05 with the following levels of significance: *p* > 0.05, non-significant, \**p* ≤ 0.05, \*\**p* < 0.01, \*\*\**p* < 0.001, and \*\*\*\**p* < 0.0001. The confidence interval in all experiments was 95% and as a detailed description of statistical parameters it is included in all figure captions. Data analysis and visualization were performed using GraphPad Prism 6.0, Fiji (ImageJ 2.1.0) Adobe Photoshop CS6 and Adobe Illustrator 25.0.

The exact sample size (n) for each experimental group/condition was given as a discrete number and unit of measurement; We provided a description of whether measurements were taken from distinct samples or whether the same sample was measured repeatedly; the statistical test(s) used AND whether they are one- or two-sided of all covariates tested; of any assumptions or corrections, such as tests of normality and adjustment for multiple comparisons; of the statistical parameters including central tendency (e.g. means) or other basic estimates (e.g. regression coefficient) AND variation (e.g. standard deviation) or associated estimates of uncertainty (e.g. confidence intervals); For null hypothesis testing, the test statistic (e.g. F, t, r) with confidence intervals, effect sizes, degrees of freedom and *P* value were noted; We gave *P* values as exact values whenever suitable.

## Supplementary figures

**Supplementary Figure 1.**
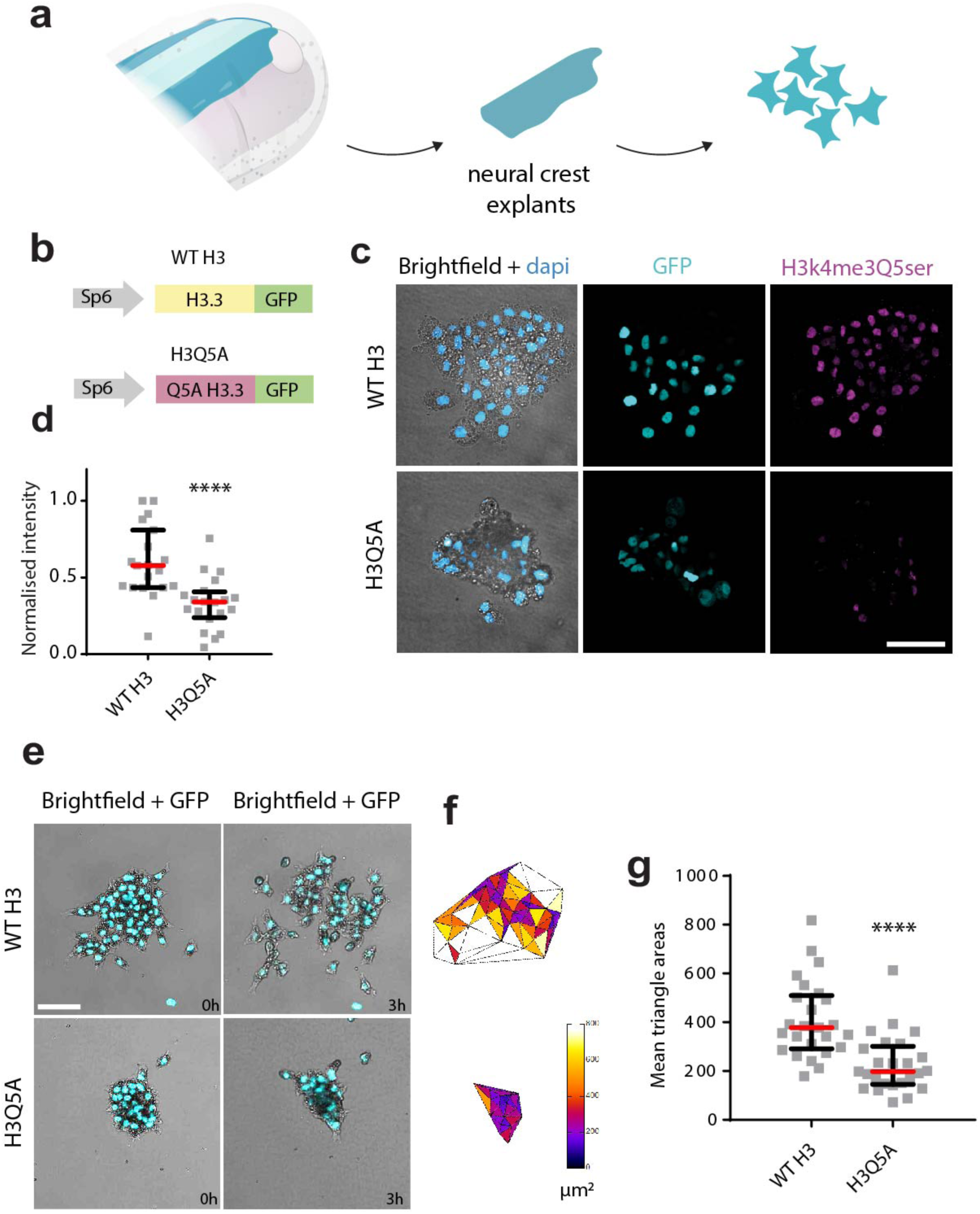
Histone serotonylation is required for neural crest migration. **a**, Schematic depicts our dispersion assay (detailed in Methods). **b**, Schematic showing WT H3 and H3Q5A constructs used to analyse the role of histone serotonylation on neural crest migration. **c,** Immunostaining for histone serotonylation in neural crest cells (magenta) after injection of WT H3 or H3Q5A (cyan) (scale bar, 50 μm). **d,** Normalised histone serotonylation fluorescence intensity. Spread of data from the indicated conditions is shown, red lines represent median and whiskers represent interquartile ranges, two-tailed Mann Whitney test **** *P*<0.0001, n_(WT H3)_= 19 explants nuclei; *n*_(H3Q5A)_= 17 explants nuclei, CI = 95%; **e-g**, neural crest dispersion analysis, related to movie S1. **e,** neural crest cells injected with WT H3 or H3Q5A (cyan) are shown 3h after plating (scale bar, 100 μm). **f, g**, Quantification of cell dispersion. **f,** Colour-coded Delaunay triangulation shown to facilitate visualization of the distances between neighbour cells. **g,** Quantification of Delaunay triangulation. Spread of data from the indicated conditions is shown, red lines represent median and whiskers represent interquartile ranges, two-tailed Mann–Whitney U-test, \*\*\*\**P* < 0.0001, n_(WT H3)_= 25 explants; n_(H3Q5A)_= 25 explants. Panels in c and e are representative examples of at least three independent experiments. CI = 95%.

**Supplementary Figure 2.**
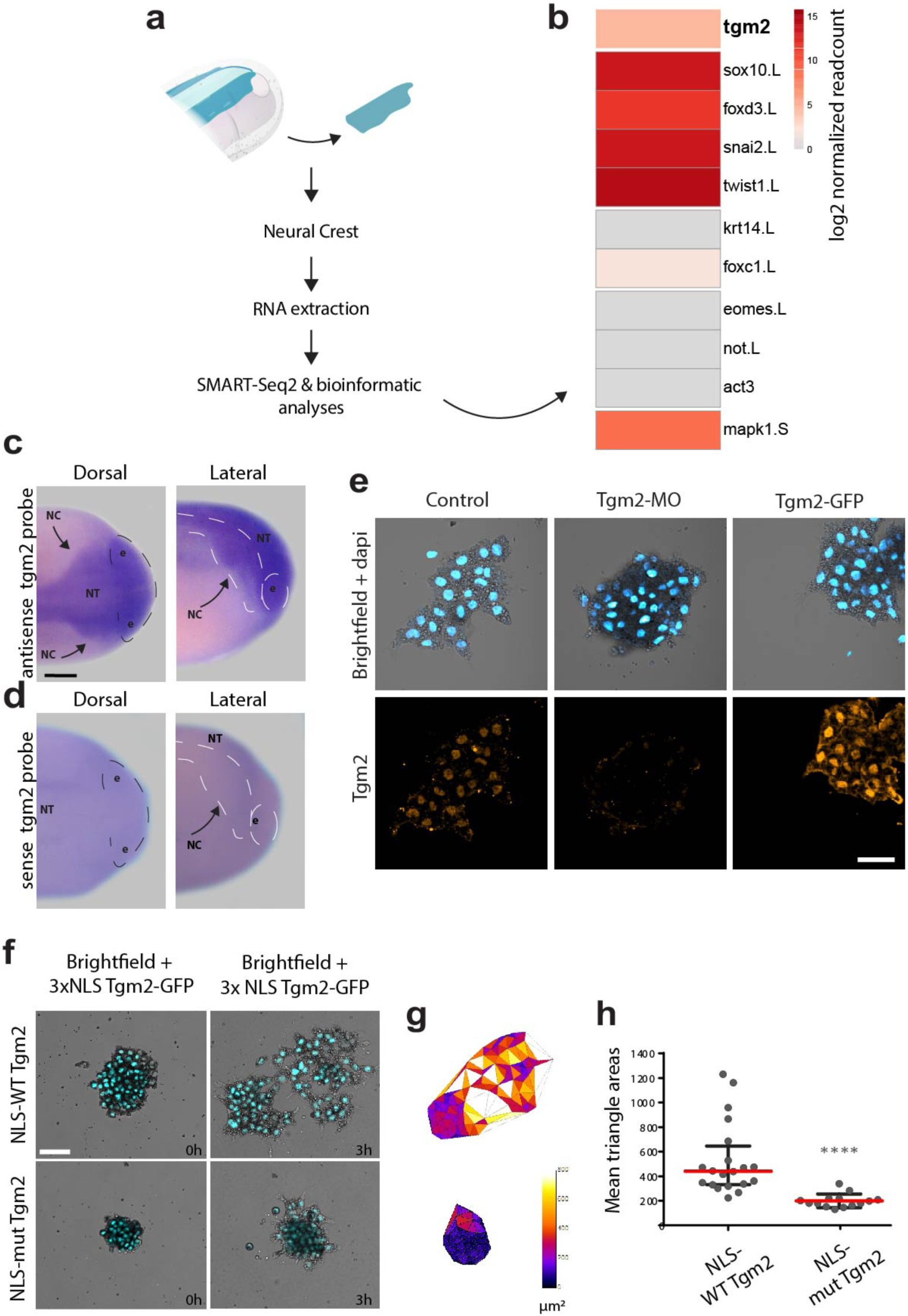
Tgm2 is expressed in the neural crest and required for migration. **a**, Schematic showing the workflow of RNA extraction from neural crest explants dissected from pre-migratory embryos at pre-migratory stage 17–18 and processed for RNA-seq. **b,** Heatmap represents log2 normalized counts confirming purity of the explants by the presence of the neural crest markers and the absence of ectodermal and mesodermal markers (ruling out contamination with surrounding tissues). **c, d**, *In situ* hybridisations against *tgm2*; **c,** antisense probe; **d,** sense probe; scale bar, 150 μm. **e,** Immunofluorescence against endogenous Tgm2 (orange) in neural crest clusters; nuclei are shown in cyan; scale bar, 50 μm; **c-e**, conditions as indicated. **f,** Neural crest cells injected with NLS-WT or NLS-mutant Tgm2 (cyan) constructs are shown 3h after plating (scale bar, 100 μm); Related to movie S2. **g, h,** Quantification of cell dispersion. **g,** Colour-coded Delaunay triangulation shown to facilitate visualization of the distances between neighbour cells. **h,** Quantification of Delaunay triangulation. Spread of data from the indicated conditions is shown, red lines represent median and whiskers represent interquartile ranges; two-tailed Mann–Whitney U-test, \*\*\*\**P* < 0.0001, n(_control_)= 20 explants; n(_Nuclear mutant_) = 15 explants. Panels in c-f are representative examples of at least three independent experiments. CI = 95%.

**Supplementary Figure 3.**
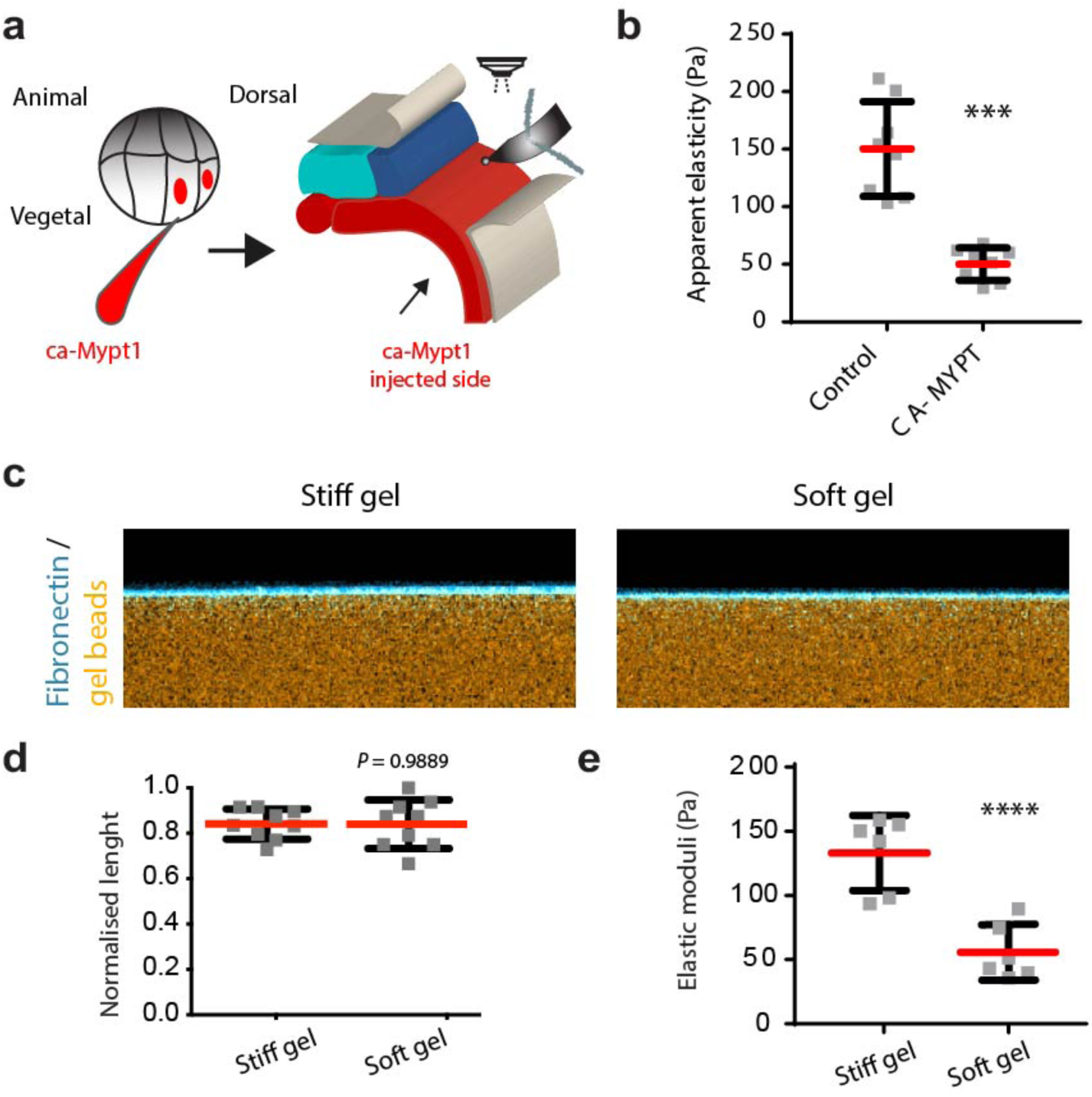
Tools to assess substrate mechanics *in vivo* and *ex vivo*. a, b, Mesoderm-targeted injections. **a,** Schematic showing that embryos were injected into two dorso-vegetal blastomeres (prospective mesoderm). **b,** iAFM measurements; spread of data is shown; red lines show mean and error bars standard deviation; two-tailed t test with Welch’s correction, \*\*\**P* = 0.0001, n_(control)_ = 8 embryos, n_(CA-MYPT)_ = 8 embryos; CI = 95%. **c-e**, Characterisation of the *ex vivo* system that reproduces the stiffness values that neural crest cells experience at non- and migratory stages. **c,** Orthogonal view of a confocal projection of soft and stiff hydrogels. Images in c are representative examples from at least 3 independent experiments. **d,** Chart showing that the layer of Fibronectin has similar thickness in soft and stiff gels; spread of data is shown; data points represent the mean obtained from each gel and at least 5 measurements were taken per gel; red line represents mean, and error bars show standard deviation (two-tailed t-test, *P* = 0.9889, CI = 95%, n = 5 gels;). **e,** AFM measurements obtained from soft and stiff hydrogels; spread of data is shown; each data point represents the mean of a gel, and 64 indentations were performed per gel; red lines show mean and error bars standard deviation; n = 5 gels; two-tailed t-test, \*\*\*\**P* < 0.0001, CI = 95%.

**Supplementary Figure 4.**
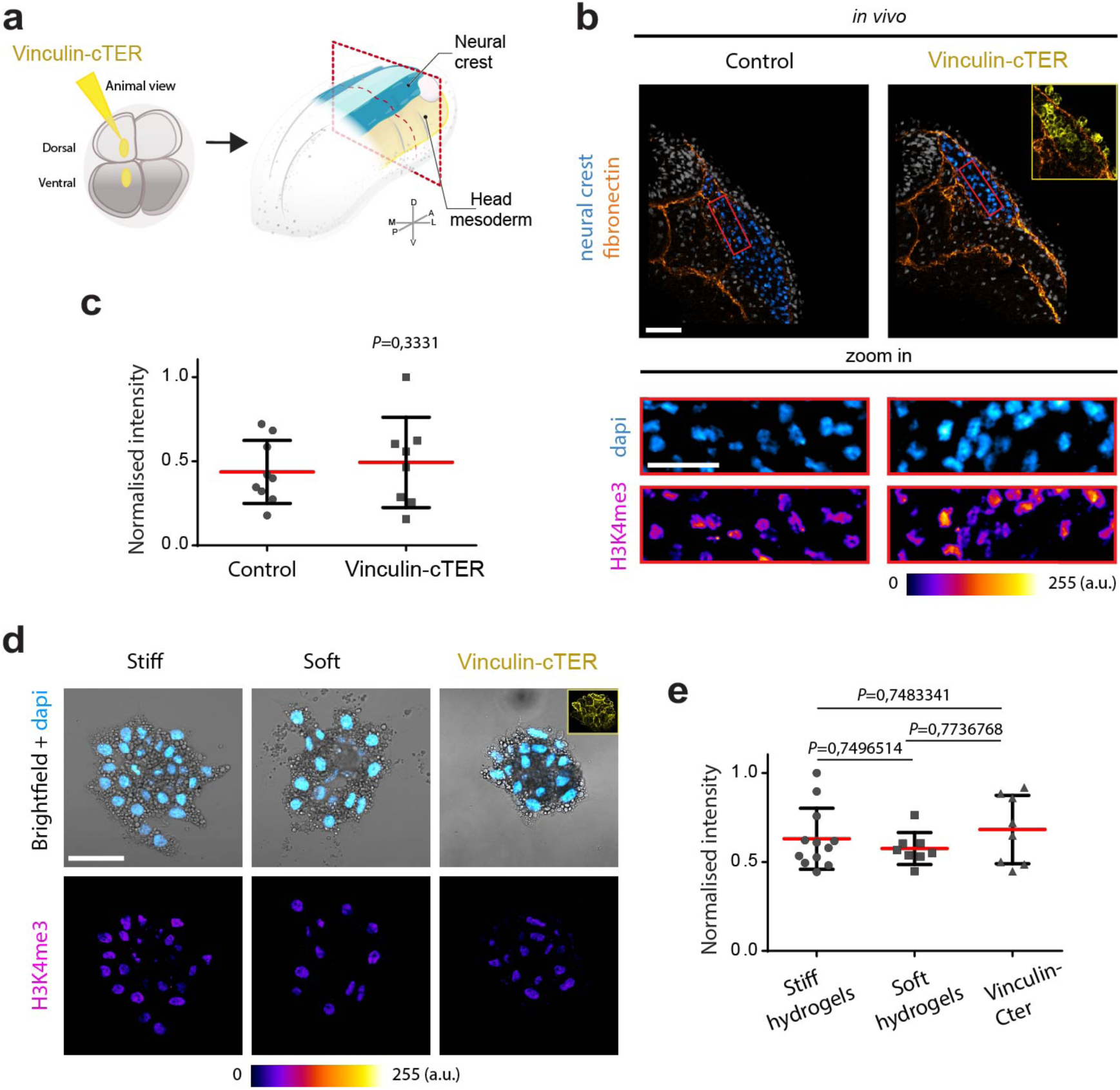
Substrate mechanics does not affect H3K4me3 levels. **a**, Schematic shows targeted injections (details in Methods) and red dashed line depicts the plane of transverse cryosection. **b,** Representative confocal projections (top-right inset shows construct expression); scale bar, 100 µm; red square indicates the region of interest shown in the lower panel (zoom-in; scale bar, 50 µm); conditions as indicated. **c,** Normalised H3k4me3 fluorescence intensity; red lines show mean and error bars the standard deviation; two-tailed t test with Welch’s correction, non-significant differences were observed, *P*= 0,3331; n_(control)_ = 9 embryos; n_(Vinculin-cTER)_ = 8 embryos. **d,** Immunostaining against H3K4me3 (colour-coded projections) in wild type neural crest clusters platted on stiff or soft hydrogels; and vinculin treated clusters (vinculin-cTER) exposed to stiff gels (yellow top-right inset shows correct injection); nuclei are shown in cyan; scale bar, 50 μm; conditions as indicated. **e,** Normalised H3k4me3 fluorescence intensity; **r**ed lines show mean, error bars represent standard deviation; ANOVA; N = 3 independent experiments; n_(stiff)_= 12 explants nuclei; *n*_(soft)_= 8 explants nuclei; *n*_(vinculin-Cter)_ = 8 explants nuclei. Panels in b and d are representative examples of at least three independent experiments. CI = 95%.

**Supplementary Figure 5.**
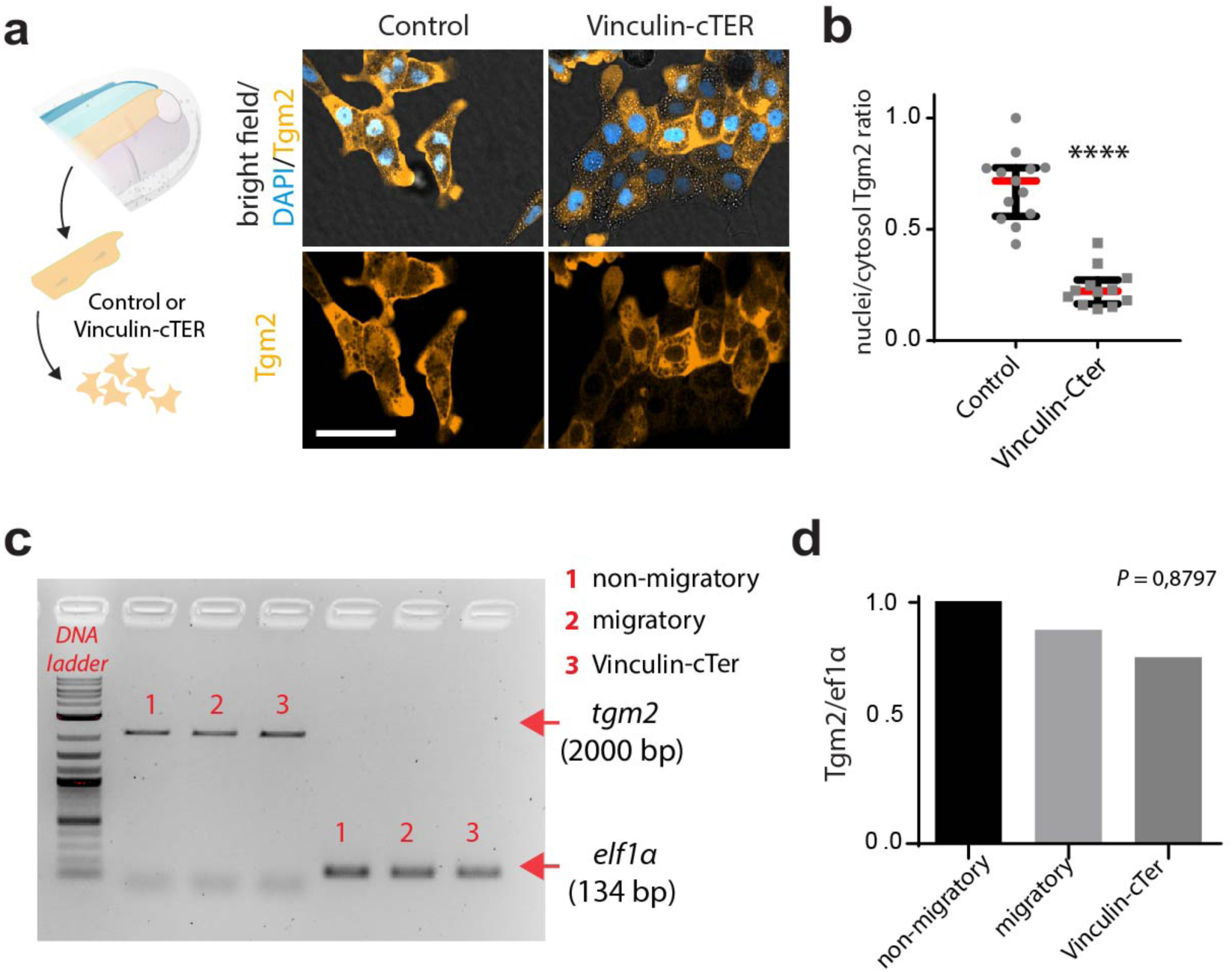
Tgm2 control at the expression and protein levels. **a**, schematic showing neural crest *ex vivo* experiments; representative confocal projections of Tgm2-GFP localization (orange); nuclei are shown in cyan; scale bar, 50 μm; conditions as indicated. **b,** Nuclear/cytosolic ratio of Tgm2-GFP fluorescence intensity; spread of data from the indicated conditions is shown, red lines represent median and whiskers represent interquartile ranges, two-tailed Mann–Whitney U-test, \*\*\*\**P* < 0.0001, n_(control)_= 13 explants nuclei; n_(Vinculin-cTer)_ = 12 nuclei. **c,** Expression of *tgm2* further assessed by semi-quantitative RT-PCR from wild type embryos along non-migratory stages (stage 13), migratory (stage 21) and vinculin-inhibited embryos (vinculin-cTER) at migratory stages. **d,** RT-PCR for *tgm2,* non-significant differences were observed, ANOVA, *P=*0,8797. Panel in a and Gel in c are representatives of 3 independent experiments.

**Supplementary Figure 6.**
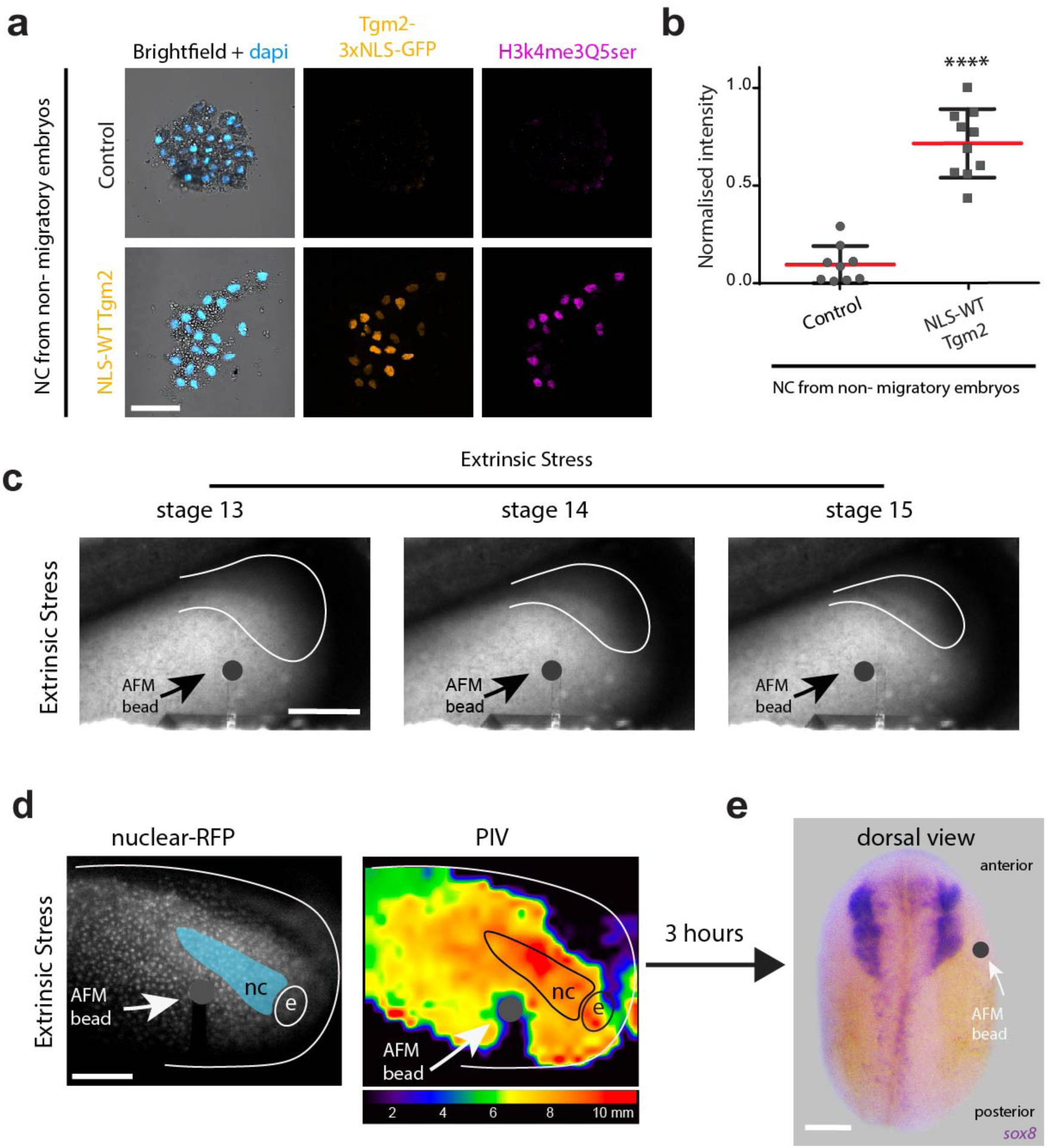
Tgm2 nuclear shuttling and mesoderm stiffening are sufficient to allow histone serotonylation and CCM. **a**, Immunostaining for histone serotonylation (magenta) in wild-type neural crest cells on non-migratory stages (stage 13), or upon injection of NLS-tagged wild-type Tgm2 (orange); nuclei are shown in cyan (scale bar, 50 μm); **b,** Normalised histone serotonylation fluorescence intensity; red lines show mean and error bars the standard deviation. Two-tailed t test with Welch’s correction, \*\*\*\**P* < 0.0001, n(non-migratory)= 9 explants; n(NLS-WT Tgm2 non-migratory)= 10 explants, CI = 95%. **c, d**, Schematic of extrinsic stress AFM experiments, images of embryos being compressed from non-migratory (stage 13) to pre-migratory stages (stage 15) with a 90-μm bead attached to an AFM cantilever (bead, black circle) are shown; scale bar, 200 μm; **d,** On the left panel, magnitude maps from a particle image velocity (PIV) analysis indicates the x–y extent of the deformation induced by AFM indentation. **e,** Dorsal view (anterior to top) of embryos hybridized with a probe against *sox8.* Panels in a and e are representatives of 3 independent experiments. CI = 95%.

**Supplementary Figure 7.**
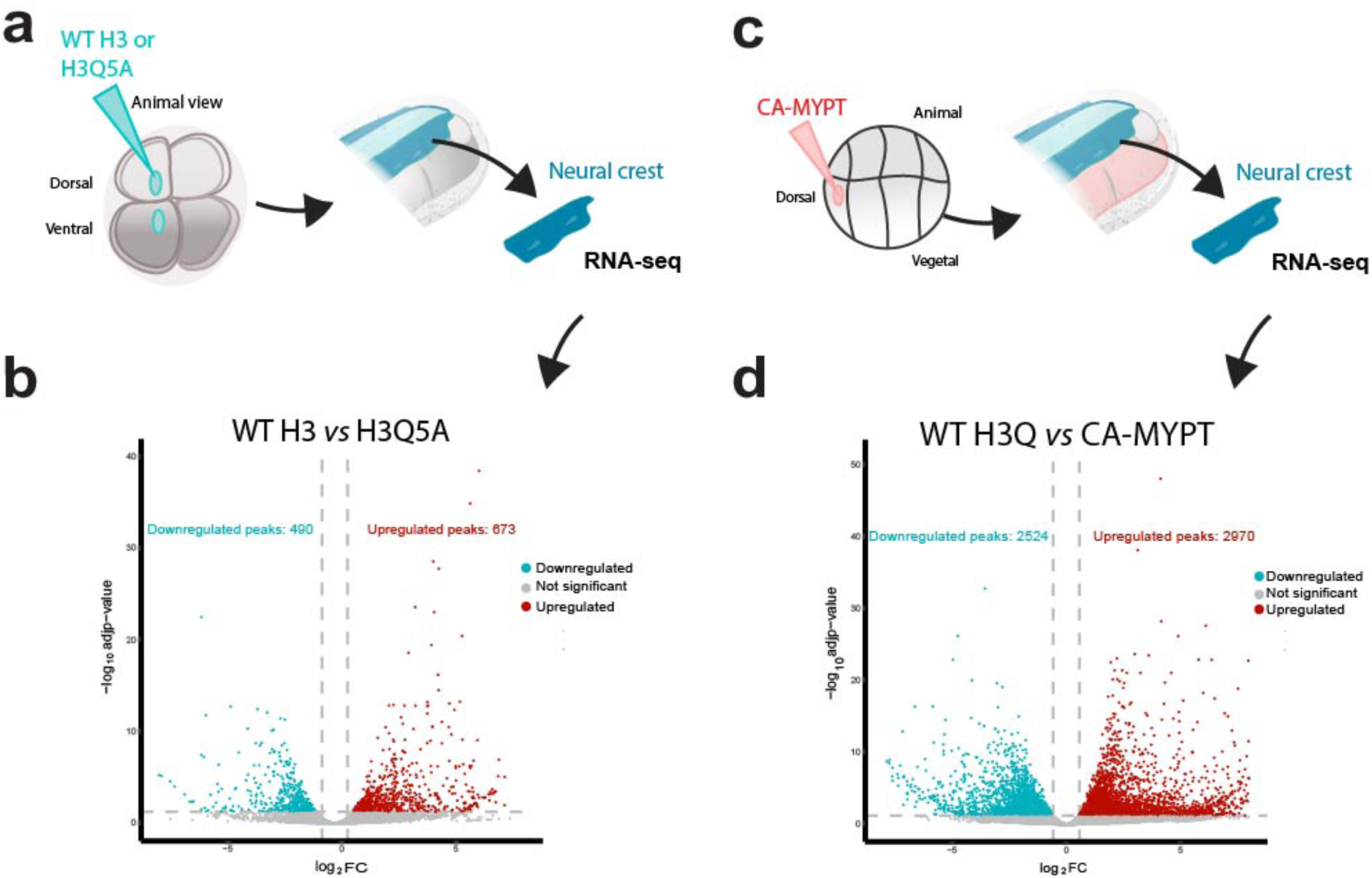
Transcriptional programme regulated by histone serotonylation in response to substrate stiffening. **a**, Schematic shows neural crest-targeted injections, dissection and RNA-seq processing (details in Methods). **b,** Volcano plot of differentially expressed transcripts controlled by histone serotonylation obtained after comparing wild-type and mutant histone H3 (H3Q5A). **c,** Schematic shows mesoderm-targeted injections, neural crest dissection and RNA-seq processing. **d,** Volcano plot of differentially expressed transcripts that rely on histone serotonylation and are downregulated by mesoderm softening (Ca-MYPT treatment). Up-regulated genes [Log2FC > 0.58, FDR-adjusted p-value < 0.05] are shown in red, down-regulated genes [Log2FC < 0.58 and FDR-adjusted p-value < 0.05] are shown in blue, and genes with no significant difference are shown in grey. Axes represent Log2FC (x-axis) and -log10 (adjusted p-value) (y-axis).

**Supplementary Table 1.**
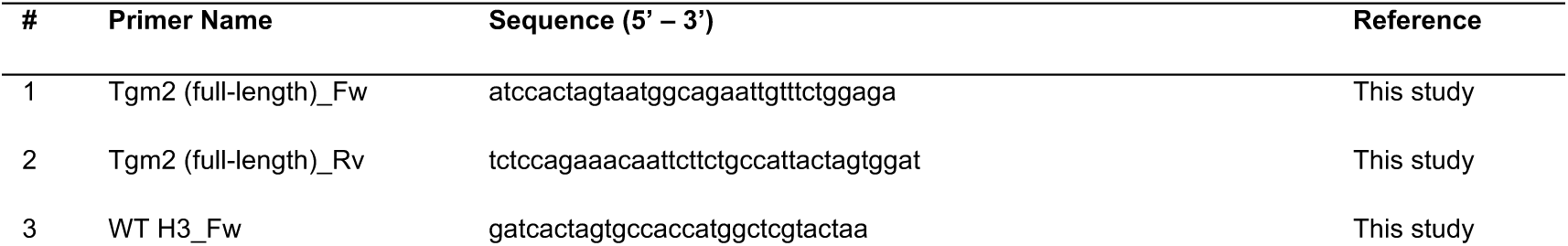
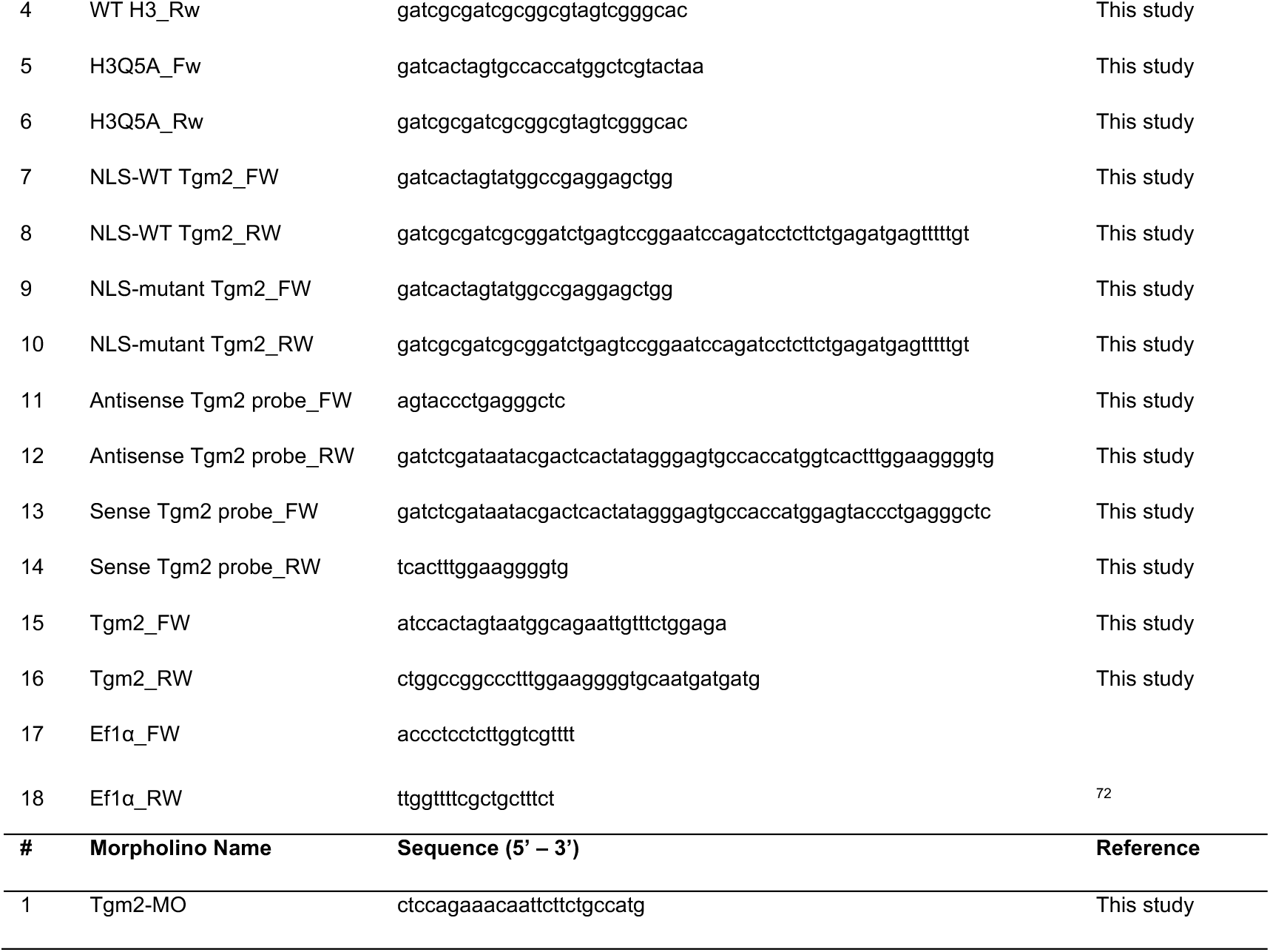
Primer and morpholino sequences used in this study.

